# Deep learning the arrow of time in brain activity: characterising brain-environment behavioural interactions in health and disease

**DOI:** 10.1101/2021.07.02.450899

**Authors:** Gustavo Deco, Yonatan Sanz Perl, Jacobo D. Sitt, Enzo Tagliazucchi, Morten L. Kringelbach

## Abstract

The complex intrinsic and extrinsic forces from the body and environment push the brain into non-equilibrium. The arrow of time, central to thermodynamics in physics, is a hallmark of non-equilibrium and serves to distinguish between reversible and non-reversible dynamics in any system. Here, we use a deep learning Temporal Evolution NETwork (TENET) framework to discover the asymmetry in the flow of events, ‘arrow of time’, in human brain signals, which provides a quantification of how the brain is driven by the interplay of the environment and internal processes. Specifically, we show in large-scale HCP neuroimaging data from a thousand participants that the levels of non-reversibility/non-equilibrium change across time and cognitive state with higher levels during tasks than when resting. The level of non-equilibrium also differentiates brain activity during the seven different cognitive tasks. Furthermore, using the large-scale UCLA neuroimaging dataset of 265 participants, we show that the TENET framework can distinguish with high specificity and sensitivity resting state in control and different neuropsychiatric diseases (schizophrenia, bipolar disorders and ADHD) with higher levels of non-equilibrium found in health. Overall, the present thermodynamics-based machine learning framework provides vital new insights into the fundamental tenets of brain dynamics for orchestrating the interactions between behaviour and brain in complex environments.

## Introduction

Survival is a key characteristic of life and requires the ability to find order in a complex, largely disordered environment. As proposed by the Austrian physicist and Nobel Laurate Ernst Schrödinger, survival is predicated on avoiding equilibrium: “How does the living organism avoid decay? … By eating, drinking, breathing and … assimilating. The technical term is metabolism” (Schrödinger, 1944). The avoidance of decay thus requires non-equilibrium interactions with the complex environment – and the brain is at the heart of these interactions.

There is a long history of understanding how the brain is able to interact with the environment. The initial extrinsic perspective proposed that the brain is primarily reflexively, driven by momentary stimulation from the environment in a *task-driven* manner (Raichle, 2010; Sherrington, 1906; Yuste *et al*., 2005). A more recent, complementary perspective was proposed by Marcus Raichle, which holds that the brain is mainly intrinsic; *resting* but switching between states whilst “interpreting, responding to, and even predicting environmental demands” (Raichle, 2006; Raichle *et al*., 2001). The evidence is clear that the brain’s metabolic energy budget for maintaining the intrinsic resting activity is much larger than that used by extrinsic task-driven demands (Zhang and Raichle, 2010). In fact, by some estimates 20% of the total energy consumption is taken up by the brain which only represents 2% of body weight (Attwell and Laughlin, 2001; Clarke and Sokoloff, 1999; Magistretti *et al*., 1999), which has led to Raichle’s poetic proposal of “dark energy” (Raichle, 2006).

The brain’s energy budget governs the flow of energy between the brain and environment, which is the ultimate cause of the non-equilibrium essential to the proposals of Schrödinger and Raichle. Any living system requires the breaking of the detailed balance of the transitions between the underlying states (Gnesotto *et al*., 2018; Schrödinger, 1944). In a system with detailed balance, the fluxes of transitions between different states disappear (Lynn *et al*., 2020; Sanz Perl *et al*., 2021). This is conveniently described in the language of thermodynamics, where a system ceases to produce entropy and becomes reversible in time (Jarzynski, 2011). In contrast, a non-equilibrium system – where the balance is broken – shows net fluxes between the underlying states, and thus becomes irreversible, establishing an arrow of time (Crooks, 1998; Feng and Crooks, 2008; Maragakis *et al*., 2008; Seif *et al*., 2021; Shirts *et al*., 2003). This is closely linked to turbulence, a classical example of non-equilibrium, which has been shown to be highly useful for optimally transferring energy/information over spacetime due to its mixing properties (Frisch, 1995). Turbulence has recently been demonstrated in the human brain, where the resulting information cascade is crucial for extracting order from disorder (Deco and Kringelbach, 2020; Sheremet *et al*., 2019).

The key idea in this paper is to quantify how the environment drives the brain by using a measure of non-equilibrium captured by the arrow of time, i.e. the level of non-reversibility in brain signals. This will allow us to assess whether the levels of non-equilibrium change in the human brain with extrinsic engagement with the environment (i.e. when performing tasks) compared to intrinsic resting state. In addition, we can assess whether the levels of non-equilibrium change with disease and therefore could be used as a potential sensitive and specific biomarker.

The ideas for the present framework comes from physics and thermodynamics, where non-equilibrium is intrinsically linked to non-reversibility (Seif *et al*., 2021) and the production of entropy, leading to the arrow of time, as originally popularised by Arthur Eddington (Eddington, 1928) and since studied in great detail (Crooks, 1998; Feng and Crooks, 2008; Jarzynski, 2011; Maragakis *et al*., 2008; Seif *et al*., 2021; Shirts *et al*., 2003). In fact, a simple, yet powerful way of assessing non-equilibrium in the brain is to directly estimate the arrow of time of the brain signals rather than the more difficult way of estimating the production of entropy (Lynn *et al*., 2020; Sanz Perl *et al*., 2021).

The non-reversibility of a physical process and the arrow of time is clearly illustrated when watching a film of a glass being shattered, which is very different from watching the same film in reverse. In thermodynamics, this can be elegantly described in terms of the entropy production, which increases when a system becomes disordered. If the total entropy production is larger than zero, it means that the system is non-reversible and in non-equilibrium. In cases such when a glass is being shattered, the non-reversibility is very clear for all to see. In contrast, a film of colliding billiards balls can be watched equally forward and backwards, making it very difficult to distinguish the correct arrow of time in the film. This process could potentially be fully reversible and not producing entropy.

In most processes, however, like the evolution of brain signals, the level of reversibility is much less clear. Here, we therefore used the power of deep machine learning to detect the level of reversibility of empirical brain imaging data. This allowed us to assess the level of non-equilibrium in brain dynamics in different states. Specifically, for whole-brain data we extracted the normal forward timeseries as well as constructing the time reversal of the backward timeseries for a given parcellation. This procedure provides a clear arrow of time for a given timeseries for which a deep learning classifier (here named Temporal Evolution NETwork, TENET) could be trained to identify the forward and the time reversal for a given timeseries of any length. The performance of TENET provides a reliable measure of the degree of non-reversibility and non-equilibrium at different levels of global and local brain organisation.

We used TENET and the concept of reversibility in thermodynamics to assess the level of non-equilibrium and arrow of time in brain dynamics in resting and seven different tasks from the large HCP neuroimaging fMRI dataset of 1003 healthy human participants. We also used the same approach to study time’s arrow in health and disease in a UCLA dataset of 265 neuropsychiatric patients (ADHD, schizophrenia and bipolar disorder as well as controls). The TENET framework turns out to be highly capable of distinguishing different tasks and different diseases and could potentially serve as a biomarker in future.

Overall, the key idea of estimating the brain’s arrow of time provides the basis for a detailed analysis of the non-stationary and non-equilibrium nature of brain dynamics in health and disease. As such, the present framework and findings provide vital new insights into the fundamental tenets of how the level of non-equilibrium of brain dynamics directly reflects the interactions with the environment, in other words how the environment drives the brain to survive and thrive.

## Results

The concept of the arrow of time is well established in physics (Jarzynski, 2011; Seif *et al*., 2021) and extensively used for problems related to thermodynamics of system in non-equilibrium including biological problems such as protein folding (Collin *et al*., 2005). In the context of neuroscience, the environment is constantly driving the brain out of equilibrium and thus the arrow of time is an excellent tool for characterising the non-equilibrium of brain signals. As a result, there has been considerable interest in using production entropy and related concepts to characterise time reversibility of brain signals (Lynn *et al*., 2020; Palus, 1996; Sanz Perl *et al*., 2021; Zanin *et al*., 2019). In contrast to these methods, here we applied Jarzynski’s idea of using a deep learning paradigm to measure the arrow of time in forward and time-reversed time series, compare the two and thus provide a direct measure of the reversibility of brain signals. We use this framework in different brain states in health and disease.

**Figures 1** and **2** provide a schematised version of the general paradigm used here. The key concept of the arrow of time is demonstrated in **Figure 1A**, which shows four sequential images from a film of a glass being shattered by a bullet. Below, the same four images are shown in a sequence in an opposite direction, i.e. in time reversal of the backward trajectory of the film. When comparing the two films, the arrow of time is very clear, which is the signature of a non-reversible physical process producing entropy in non-equilibrium. More general, as shown in **Figure 1B**, the field of thermodynamics in physics can be used to describe such processes associated with the production of entropy and consequently with non-equilibrium. The figure shows the evolution over time of a non-equilibrium system with two states A and B and their associated trajectories. The forward and backward trajectories of the movies in **Figure 1A** are described as forward (A → B, black arrow) and backward (B → A, red arrow) processes. The time reversal of the backward trajectory (red stippled arrow) can be thought of as the movie of the backward trajectory that is played forward in time (see bottom of **Figure 1A**). A non-reversible process results from when the ability to differentiate between the trajectories in time described by the forward (black arrow) and time reversal (stippled red arrow). The second law of thermodynamics (usually attributed to Rudolph Clausius and Sadi Carnot) states that if the entropy production is larger than zero, this corresponds to non-reversibility of a non-equilibrium system. In contrast, if there is no entropy production, this describes a reversible, equilibrium system.

**Figure 1.**
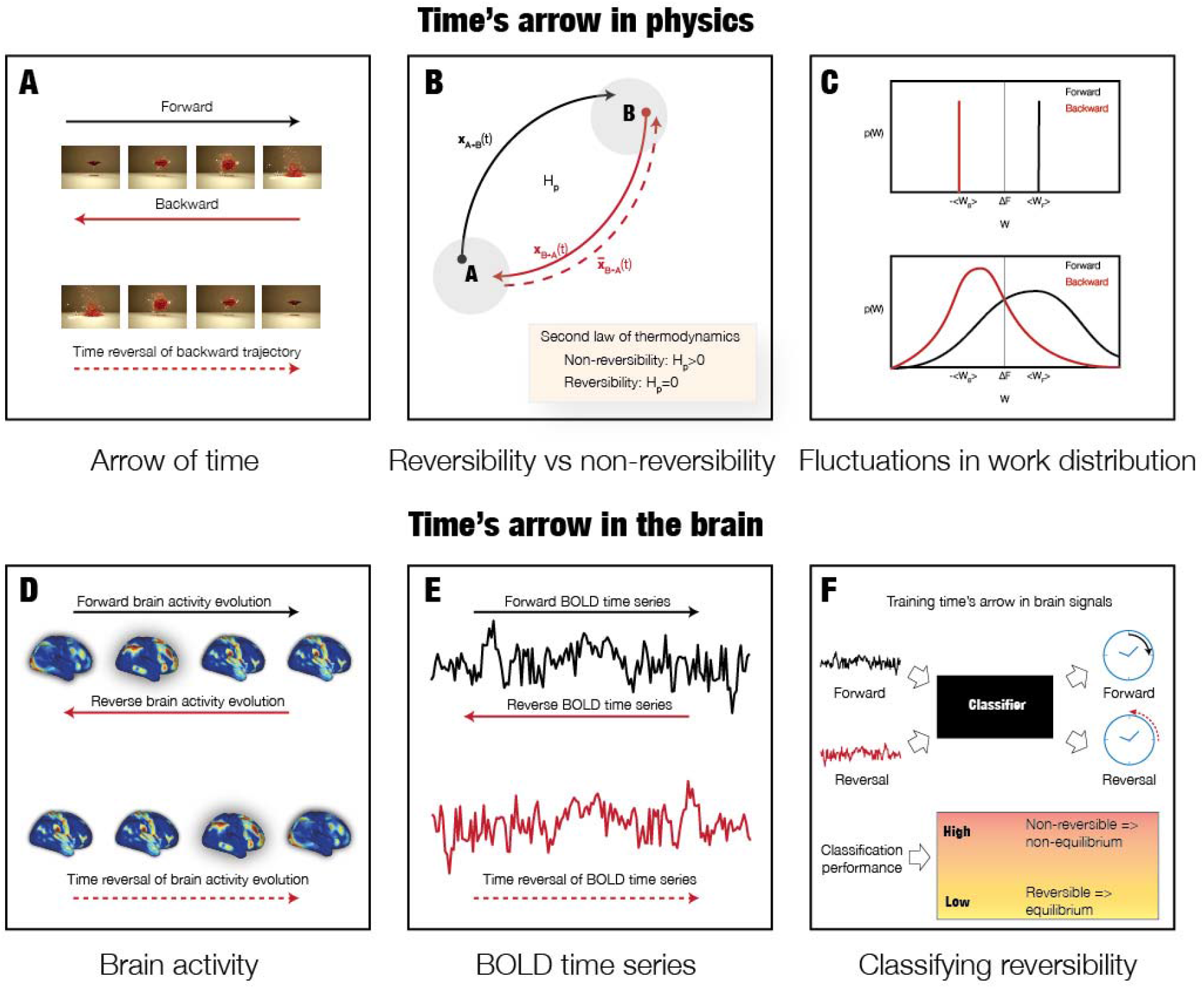
The arrow of time in physics and brain dynamics. **A)** The sequence of the four top images shows a glass being shattered by a bullet and we clearly perceive the causal passage of time, also called the arrow of time. In contrast, this cause and effect is shattered by showing these images backward - by time reversing the backward evolution. This means that this process is non-reversible. **B)** In thermodynamics, non-reversibility can be associated with the production of entropy. The figure shows a non-equilibrium system with two states A and B and the associated trajectories evolving during forward (A → B, black arrow) and backward (B → A, red arrow) processes. Both the forward and backward trajectories can be depicted as the movie shown in the top of panel A – but with a different arrow of time. In contrast, the time reversal of the backward trajectory (red stippled arrow) can be imagined as the movie of the backward trajectory that is played forward in time (see bottom of panel A). If the forward and time reversal of the backward trajectories are different, this corresponds to non-reversibility of the process. The second law of thermodynamics uses the entropy production to describe this. If the entropy production is larger than zero, this corresponds to non-reversibility of a non-equilibrium system. In contrast, if there is no entropy production, this is a reversible, equilibrium system. **C)** More specifically, when small systems undergo thermodynamic processes, the fluctuations are non-negligible and the second law of thermodynamics expresses this in terms of averages. The so-called Clausius inequality establishes that the work W (averaged over many repetitions) is larger than the change in its free energy ΔF. The figure shows the work distribution p(W) for the average of the work associated with the forward and backward trajectories, denoted <W_F_> and <W_B_>, respectively. For non-reversible macroscopic processes (like the movie shown in panel A) fluctuations are negligible and the distinction is clear between the distribution of work (top of panel) and therefore arrow of time is easy to establish. In contrast, in microscopic systems (such as brain signals) the average work is similar but the fluctuations are more pronounced and therefore the differences in distribution less clear. In this case it is much harder to establish the arrow of time, and consequently establish whether the system is non-equilibrium and non-reversible. Therefore, to solve this problem, we used deep learning in empirical brain imaging data to detect the reversibility of the system (see Figure 2). **D)** First, we used large-scale empirical whole-brain neuroimaging data from over 1000 participants when resting and performing seven different tasks. **E)** From this data, we were able to extract the forward timeseries as well as constructing the time reversal of the backward timeseries for a given parcellation. **F)** This procedure provides a clear arrow of time for a given timeseries and allow us to train a classifier to identify the forward and the time reversal for a given timeseries of any length. The classification performance provides a measure of the degree of non-reversibility and non-equilibrium.

**Figure 2.**
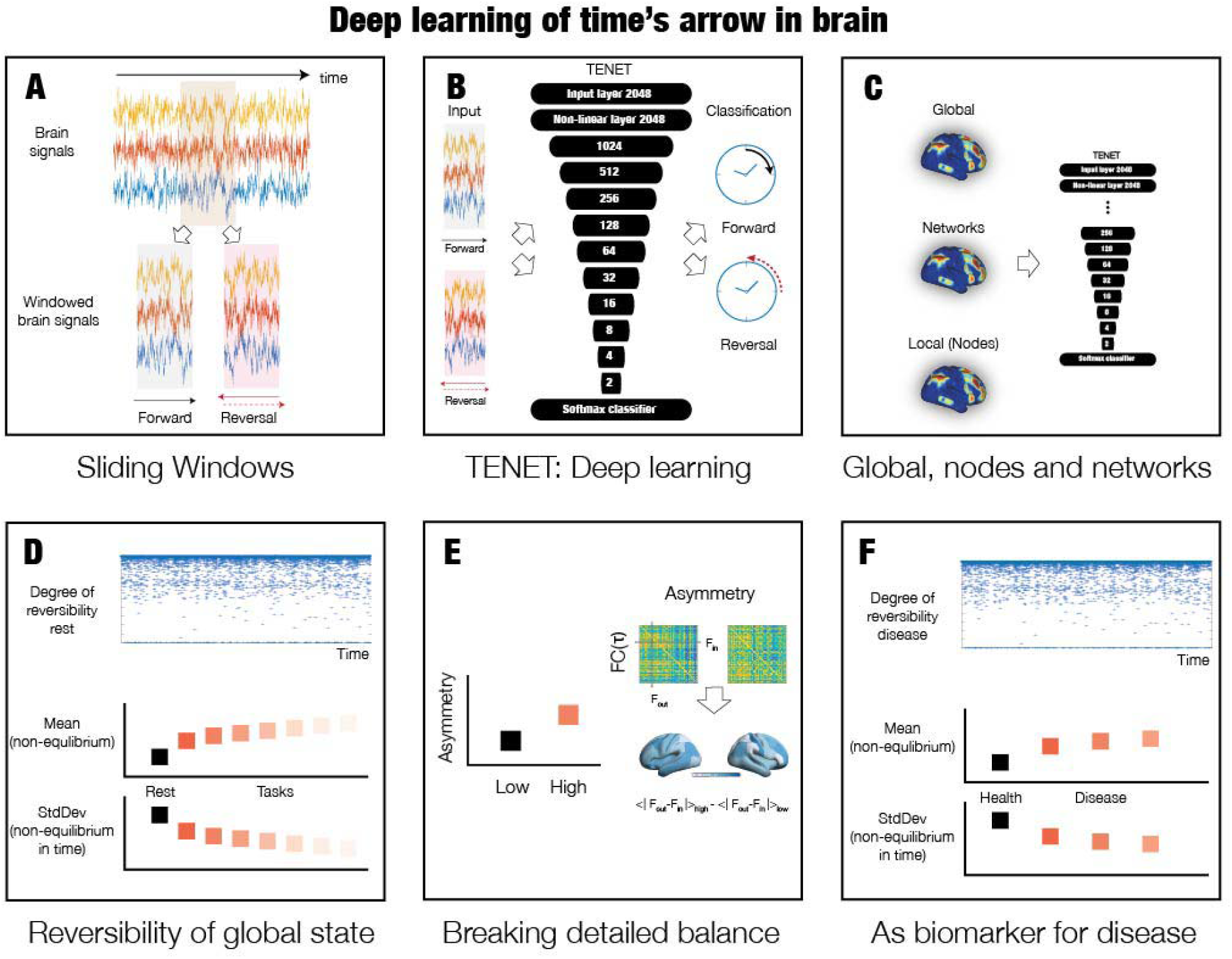
Deep learning the arrow of time in health and disease. In order to discover the arrow of time in brain dynamics in health and disease, we designed a deep learning pipeline named Temporal Evolution NETwork (TENET). **A)** Specifically, we used sliding windows of brain signal timeseries from all brain regions in all participants. **B)** These sliding windows were then used in the TENET, a deep learning network classifier with 13 layers for classifying the arrow of time. **C)** This strategy allowed us to study non-reversibility and non-equilibrium at different levels of granularity, from global (all signals) to network level to individual node level signals. **D)** After training, TENET was able to characterise the degree of reversibility, i.e. non-equilibrium for each sliding window (top panel). We performed this procedure on data resting and seven tasks and computed the means of the levels of certainty of the classifier output (across time) as a measure of the degree of non-equilibrium (middle panel). The standard deviation of this measure establishes stability of this non-equilibrium across time. Given that non-equilibrium states are already non-stationary, this provides the second order of non-stationarity (see Methods). **E)** Non-equilibrium is associated with the breaking of detailed balance of a system. We estimated this by selecting windows of low and high reversibility, and computing the FC(τ), i.e. the time-delayed functional connectivity between all pairs of brain regions. Specifically, the degree of asymmetry of the FC(τ) matrix is a proxy for the breaking of the detailed balance with more asymmetry corresponding to more unbalance. The level of asymmetry can also be rendered on the brain (see Methods). **F)** Finally, we used TENET on resting state data from neuropsychiatric patients with diagnoses of schizophrenia, ADHD, and bipolar disorder, as well as age-matched controls. Different levels of non-equilibrium provide a potential biomarker of neuropsychiatric disease.

In thermodynamics, the Clausius inequality establishes that the work W associated with the process (averaged over many repetitions) is larger than the change in its free energy ΔF. **Figure 1C** shows distributions of the work p(W) for the average of the work associated with the forward and backward trajectories, denoted <W_F_> and <W_B_>, respectively. For non-reversible macroscopic processes (like the movie shown in **Figure 1A**) fluctuations are negligible and the distinction is clear between the distribution of work (top of panel) and therefore the arrow of time is easy to establish. In contrast, in microscopic systems (which includes brain signals) the average work is similar, but the fluctuations are more pronounced and therefore the differences in distribution less clear. In such cases it is much harder to establish the arrow of time, and thus establish whether a system is non-equilibrium and non-reversible.

This uncertainty is a perfect case for which to use advanced machine learning techniques. Here, we used deep learning in empirical brain imaging data to detect the reversibility of the system. **Figure 1D** and **Figure 1E** illustrate how we used whole-brain activity from large-scale empirical whole-brain neuroimaging data from over 1000 participants to construct the forward and time-reversed timeseries needed to establish the arrow of time and hence non-equilibrium by detecting the level of non-reversibility. Specifically, **Figure 1F** illustrates how we extracted the forward timeseries as well as constructing the time reversal of the backward timeseries for the DK80 parcellation (see Methods). This procedure establishes a clear arrow of time for any given timeseries, which was used to train a classifier to identify the forward and the time reversal. If the classification performance is high, this provides evidence for non-reversibility and non-equilibrium, while low performance implies the opposite.

**Figure 2** specifies the full learning pipeline using a deep learning Temporal Evolution NETwork (TENET) to establish the arrow of time. **Figure 2A** shows how we used sliding windows of brain signal timeseries from all brain regions in all participants. **Figure 2B** shows how these sliding windows were then used in TENET, a deep learning network classifier with 13 layers for classifying the arrow of time. **Figure 2C** shows how this strategy allowed us to study non-reversibility and non-equilibrium at different levels of granularity, from global (all signals) to network level to individual node level signals.

**Figure 2D** shows how TENET should be able to characterise the degree of reversibility, i.e. non-equilibrium for each sliding window (top panel). We trained the TENET on a large dataset of data resting and seven tasks and, on a validation dataset, computed the means of the levels of certainty of the classifier output (across time) as a measure of the degree of non-equilibrium (middle panel). The standard deviation of this measure establishes stability of this non-equilibrium across time. Given that non-equilibrium states are already non-stationary, this provides the second order of non-stationarity (see Methods). **Figure 2E** shows that non-equilibrium is associated with the breaking of detailed balance of a system. We estimated this by selecting windows of low and high reversibility, and computing the FC(τ), i.e. the time-delayed functional connectivity between all pairs of brain regions. Specifically, the degree of asymmetry of the FC(τ) matrix is a proxy for the breaking of the detailed balance with more asymmetry corresponding to more unbalance. Finally, **Figure 2F** shows how TENET can be used on resting state data from neuropsychiatric patients with diagnoses of schizophrenia, ADHD, and bipolar disorder, as well as age-matched controls. Computing the different levels of non-equilibrium provide a potential biomarker of neuropsychiatric disease.

### Significant differences in global levels of non-equilibrium/non-reversibility in HCP rest and seven tasks

For the global level of analysis of how the environment is driving the brain out of equilibrium, we extracted BOLD time series from the DK80 parcellation covering the whole brain in rest and the seven tasks. For each of HCP participant, we extracted forward and backward patterns in sliding windows with a length of 20 TRs (of 0.72sec), which were then shifted 3 TRs forward. Each of the sliding windows consisted of two input patterns containing 1) forward and 2) time reversed backward sliding windowed timeseries, which was each labelled with an output class label of forward and backward, respectively.

For the training of TENET, in order to perfectly balance the data and avoid any potential source of bias, we used 890 HCP participants with the longest possible duration available in all conditions (176 TRs). For generalisation, we performed the data analysis on a separate 100 HCP participants. The data analysis consisted of computing the level of non-equilibrium/non-reversibility, R(t), using the output of TENET on this generalisation set after being trained on the bulk of the data.

As specified in details in the Methods, R(t) is computed as the accuracy of classification of forward and time reversal of backward trajectory of the global timeseries (across sliding windows at time t and across participants). Perfect classification of maximal non-reversibility is thus assigned a value of 1 and where 0 corresponds to full reversibility.

**Figure 3A** (left panel) contains a boxplot showing that the brain dynamics during REST have significantly lower levels of reversibility than in tasks (p<0.001, Wilcoxon rank sum). As can be seen, the highest level of non-equilibrium/non-reversibility is found in the SOCIAL task, reflecting how the environment is forcing a stronger arrow of time and thus non-reversibility. But, equally, the other tasks, ordered by levels of non-equilibrium/non-reversibility (RELATIONAL, EMOTION, GAMBLING, MOTOR, WM (working memory) and LANGUAGE) are significantly more driven by the environment than during REST. It is interesting to note the significant differences between the tasks too (p<0.001, Wilcoxon rank sum; all significant comparisons except for MOTOR vs LANGUAGE and WM and GAMBLING).

**Figure 3.**
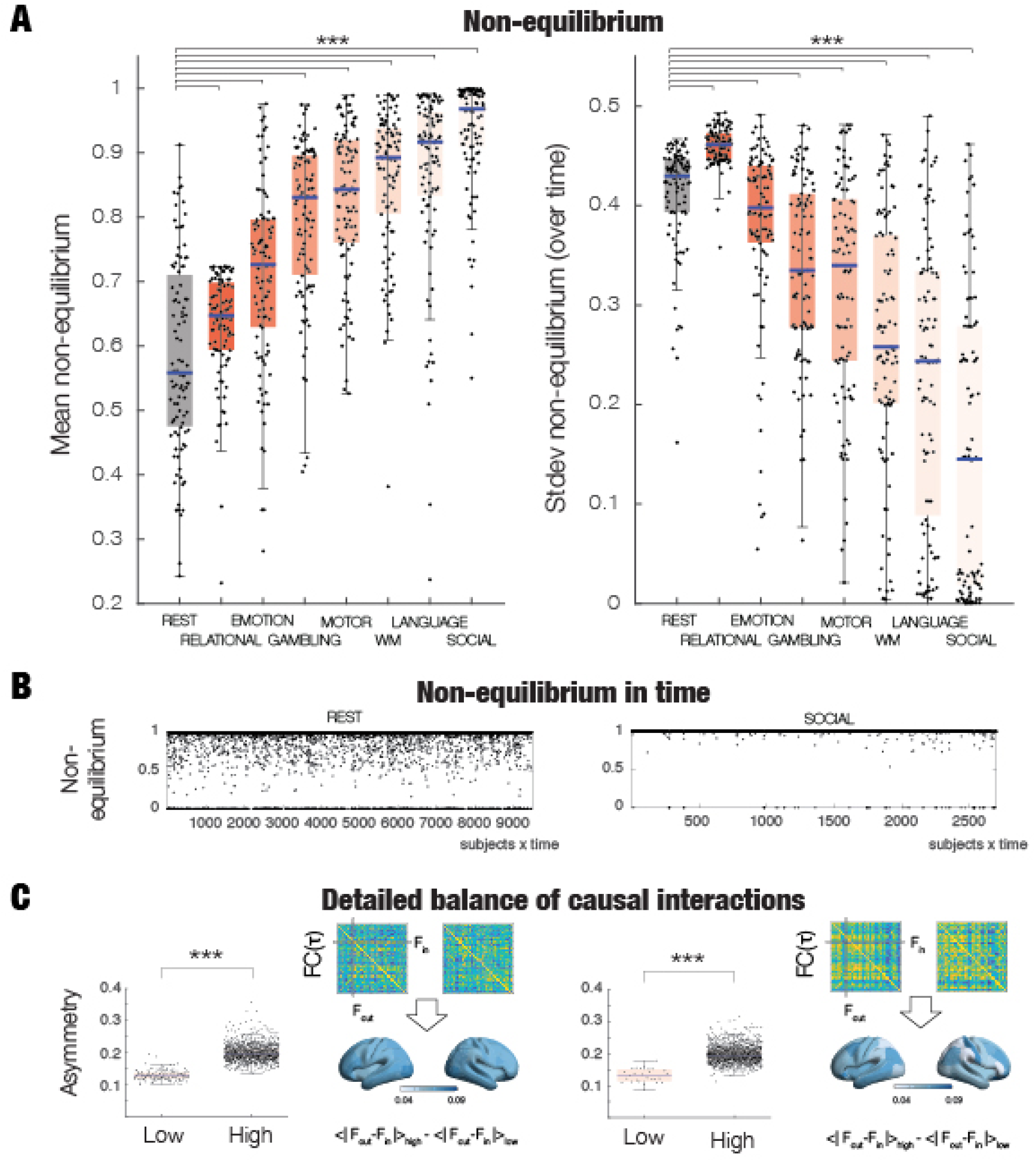
Global non-equilibrium in HCP rest and seven tasks. **A)** Left panel shows the mean non-equilibrium for rest and the seven tasks ordered by the increase in their mean level of non-equilibrium. The level of non-equilibrium/non-reversibility, R(t), is computed using the output of TENET on a 10% validation set after being trained on 90% of the data. In brief, R(t) is computed as the accuracy of classification of forward and time reversal of backward trajectory of global timeseries (across sliding windows at time t), where a value of 1 corresponds to perfect classification, i.e. maximal non-reversibility (see Methods). As can be seen from the boxplot, brain dynamics during rest exhibits significantly lower levels of reversibility than that found in tasks (p<0.001). The highest level of non-equilibrium/non-reversibility is found in the Social task, reflecting a stronger arrow of time associated with an increase in the interaction with the environment. In other words, the brain dynamics in tasks are showing more non-equilibrium than rest and are thus more driven by the environment. The right panel shows the stability of this non-equilibrium across time, i.e. providing a measure of second order of non-stationarity. Brain activity during rest is showing significantly more variability in the second order of non-stationarity than tasks (p<0.001). **B)** The panel shows the level of non-equilibrium/non-reversibility, R(t), over time for rest (left panel) and the social task (right panel). Note how the evolution of R(t) is more variable in rest. **C)** Causal interactions vary across time and consistently show a significantly stronger breaking of the detailed balance in windows with high compared to low levels of non-equilibrium/non-reversibility (compare low and high boxplots, p<0.001) for both rest (left panel) and the social task (right panel). This is measured as the asymmetry of the time-shifted functional connectivity (see Methods). The renderings of brains reflect which brain regions are showing more symmetry-breaking between low and high levels of non-equilibrium/non-reversibility. The brain shows more heterogenous patterns of change during the social task than in rest (compare right with left panel). This strongly suggests that when performing a complex task, the environment is driving the brain in very specific ways to a more non-equilibrium state.

In addition, the right panel shows the level of non-stationarity, which is the standard deviation of the levels of non-equilibrium across time. The differences between rest and tasks were similar to the results of the mean of the non-equilibrium in the sense that there were significant differences between all (p<0.001, Wilcoxon rank sum; except between MOTOR vs LANGUAGE and WM and GAMBLING) but importantly for this measure, the SOCIAL task had the lowest variability over time, which was much lower than REST. On the other hand, REST is showing one of the largest levels of non-stationarity which is consistent with the idea that resting state is more intrinsically driven and less extrinsically controlled by the environment. This can also be appreciated from **Figure 3B**, where the two panels show time evolution of the levels of non-equilibrium/non-reversibility, R(t), for REST (left) and the SOCIAL task (right).

As mentioned above, the level of equilibrium is associated with the fluxes of transitions between different states, i.e. how the detailed balance of the transitions between the underlying states disappear in completely equilibrium. In thermodynamics, a non-equilibrium system contains net fluxes between the states as a function of broken balance, which is the source of non-reversibility and thus the arrow of time (Crooks, 1998; Feng and Crooks, 2008; Maragakis *et al*., 2008; Seif *et al*., 2021; Shirts *et al*., 2003). In order to establish a direct link between our measure of non-equilibrium/non-reversibility and broken detailed balance, we measured the asymmetry of the time-shifted functional connectivity (see Methods).

In brief, in order to measure the causal interactions, we selected patterns from sliding windows of low and high reversibility, and computed the time-delayed functional connectivity matrix, *FC*(τ), between all pairs of brain regions, over all participants and all sliding windows for each condition of HCP REST and the SOCIAL task, which is the task with most non-equilibrium. The global level of asymmetry was computed for each sliding window as the mean value over the elements of the difference between this matrix and its transposed. In contrast, for the node-level of asymmetry, we first computed the incoming and outgoing regional flow for each sliding window and then computed the average over all sliding windows and participants of the absolute difference between the two regional flows (see Methods for detailed information). We render the change between high and low levels of the node-level asymmetry.

As can be seen in **Figure 3C**, we found significantly stronger breaking of the detailed balance in windows with high compared to low levels of non-equilibrium/non-reversibility (compare low and high boxplots, p<0.001, Wilcoxon rank sum). On the right of the boxplot, we show an example of the asymmetry matrices for a single participant at a given timepoint. Below renderings are shown of the change between low and high levels of the node-level asymmetry.

Consistent with the close link between symmetry-breaking and our measure, we found more heterogenous patterns of change during the SOCIAL task than in REST. This again demonstrates that when engaged in a task, the environment is driving the brain in very specific ways to higher levels of non-equilibrium.

### Network-level analysis of HCP data differentiates between HCP and task

We performed a network level analysis of HCP data using the TENET framework. For this analysis we used the Schaefer 500 parcellation where each parcel belongs to one of the seven Yeo networks. The TENET framework was then applied in the same manner as in the global analysis but now used on the parcels belonging to each network. Again, in order to balance the data, we used 890 HCP participants with the longest possible duration available in all conditions (176 TRs). The results are from the generalisation that was performed on a separate 100 HCP participants (see Methods). We used the same sliding window size and shifting of this window as in the global level analysis but now the input is the window size multiplied by the number of parcels for a given level analysis of HCP rest and tasks. Analysis of the non-equilibrium/non-reversibility of the seven Yeo resting state networks showed differential responses between rest and tasks for the seven resting state networks.

**Figure 4A** shows boxplots of the level of non-equilibrium for each of the seven Yeo networks for rest and the seven tasks, which is also shown as spider plots in **Figure 4B**. Comparable to the global level analysis there is lower levels of non-equilibrium for rest compared to the seven tasks. This suggests that REST is more in equilibrium and therefore more intrinsically driven. Interestingly, the REST condition shows the highest levels of non-equilibrium in the default mode network (DMN) and Visual (VIS) network. Similarly, the sensory networks (VIS and SOM) are showing the highest level of non-equilibrium across the seven tasks (except for LANGUAGE), and thus most driven by the environment. The lowest levels of non-equilibrium are found in the Limbic network (LIM), signalling that the LIM network is more intrinsically driven. Of the seven tasks the lowest levels of non-equilibrium are found in the EMOTION and RELATIONAL tasks, which are almost as low as REST.

**Figure 4.**
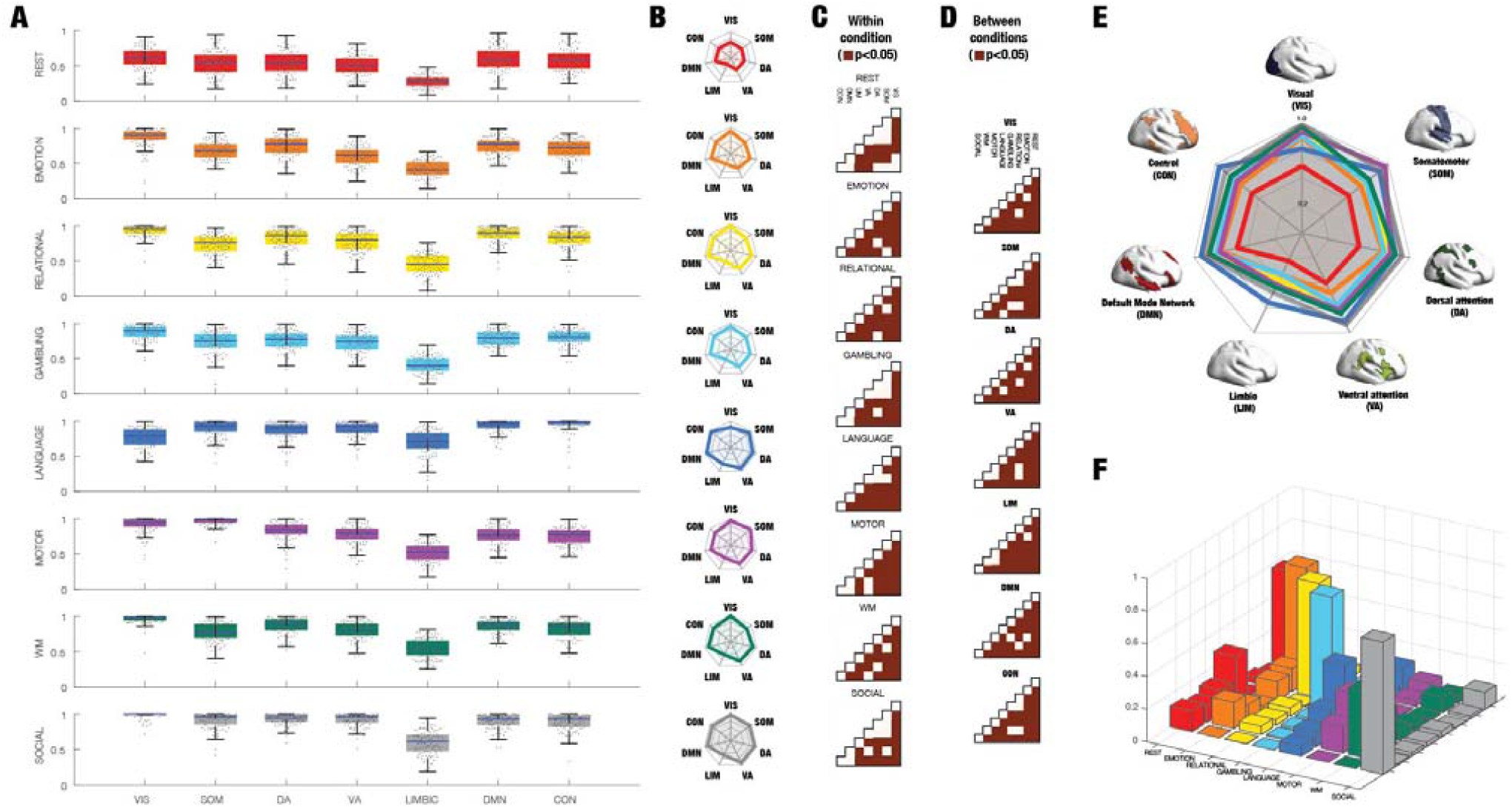
TENET network level analysis of HCP rest and tasks. Analysis of the non-equilibrium/non-reversibility of the seven Yeo resting state networks showed differential responses between rest and tasks for the seven resting state networks. **A)** Boxplots of the level of non-equilibrium for each of the seven Yeo networks for rest and the seven tasks. Similar to the global level analysis there is lower levels of non-equilibrium for rest compared to the seven tasks, suggesting, as expected, that REST is more intrinsically driven and thus more in equilibrium. It is of interest to note that REST is characterised by having the highest levels of non-equilibrium in the default mode network (DMN) and Visual (VIS) network. Equally, across the tasks, except for LANGUAGE, the sensory networks (VIS and SOM) show the highest level of non-equilibrium and thus most driven by the environment. Interestingly, again except for LANGUAGE, the Limbic network (LIM) exhibit the lowest levels of non-equilibrium, signalling that it is more intrinsically driven. Overall, of the seven tasks, the Yeo networks in EMOTION and RELATIONAL show almost as low levels of non-equilibrium levels as REST. **B)** Corresponding spider plots of the level of non-equilibrium for each Yeo network in rest and seven tasks, which are colour coded as in panel A. **C)** Within condition (rest and the seven tasks) significance testing between the level of non-equilibrium of the seven Yeo networks is shown in the upper quadrangle of the matrices (with brown squares signifying p<0.05) Almost all comparisons within conditions are significant but less so for REST. **D)** Similar, the matrices show between conditions significance testing between the level of non-equilibrium of the seven Yeo networks is shown in the upper quadrangle of the matrices (with brown squares signifying p<0.05). Almost all comparisons across conditions are significant. **E)** To facilitate comparisons, we show a combined spider plot of the level of non-equilibrium for each Yeo network in rest and seven tasks (with the Yeo networks rendered on the brain with a separate colour coding). **F)** Classification of conditions (rest and seven tasks) using the network level TENET output. As can be seen from the confusion matrix, the SVM provides excellent classification results much above chance level. Average classification for the diagonal is 59% with a chance level of 12.5%.

The differences across conditions and across networks are quantified *within* conditions in **Figure 4C**, showing significant differences (p<0.05, Wilcoxon rank sum) and *between* conditions in **Figure 4D**. As can be seen most of the comparisons are significant, suggesting that the specificity of level of non-equilibrium/non-reversibility within and between conditions. **Figure 4E** further visualises these differences between conditions by showing a combined spider plot of the levels of non-equilibrium for each Yeo network in all conditions (rest and seven tasks).

We checked the possibility of classifying the conditions based on the network level TENET output. Using a support vector machine (SVM) with Gaussian kernels on the 100 HCP participants used for generalisation. For the SVM, we subdivided the 100 participants into 90% training and 10% validation, repeated and shuffled 100 times. The SVM had seven inputs (the Yeo resting state networks) corresponding to the output produced by the network level TENET. The output was eight classes corresponding to the conditions (rest and seven tasks). **Figure 4F** shows the resulting confusion matrix, which provides excellent classification results much above chance level with an average classification accuracy of 59% (on the diagonal) compared with the chance level of 12.5%. Interestingly, the results of classifying rest vs all tasks, produced a very high accuracy of 93.1% on the generalisation dataset, using exactly the same procedure as for classifying the individual tasks.

### Node-level analysis of HCP data reveals distinct patterns of local non-equilibrium/non-reversibility

Beyond the global and network level analyses, we were interested in studying how the node-level non-equilibrium/non-reversibility can distinguish the local level of influence on brain regions by the environment.

We applied the TENET framework at the node-level using exactly the same amount of data across all rest and seven tasks, similar as above (see Methods). For the training of TENET, in order to perfectly balance the data and avoid any potential source of bias, we used 890 HCP participants with the longest possible duration available in all conditions (176 TRs). For generalisation, we performed the data analysis on a separate 100 HCP participants.

The node-level rendering for non-equilibrium/non-reversibility for REST and the SOCIAL task is shown in **Figure 5A**. Similar to the global results, these are the two conditions with the lowest and highest levels of non-equilibrium (compare lighter shades of brown for REST to the darker for SOCIAL). However, here we were also able to draw out the inter-regional heterogeneity. To further draw out the differences between tasks, in **Figure 5B**, we render the thresholded node-level of non-equilibrium for all the seven tasks (thresholded to include the top 15%).

**Figure 5.**
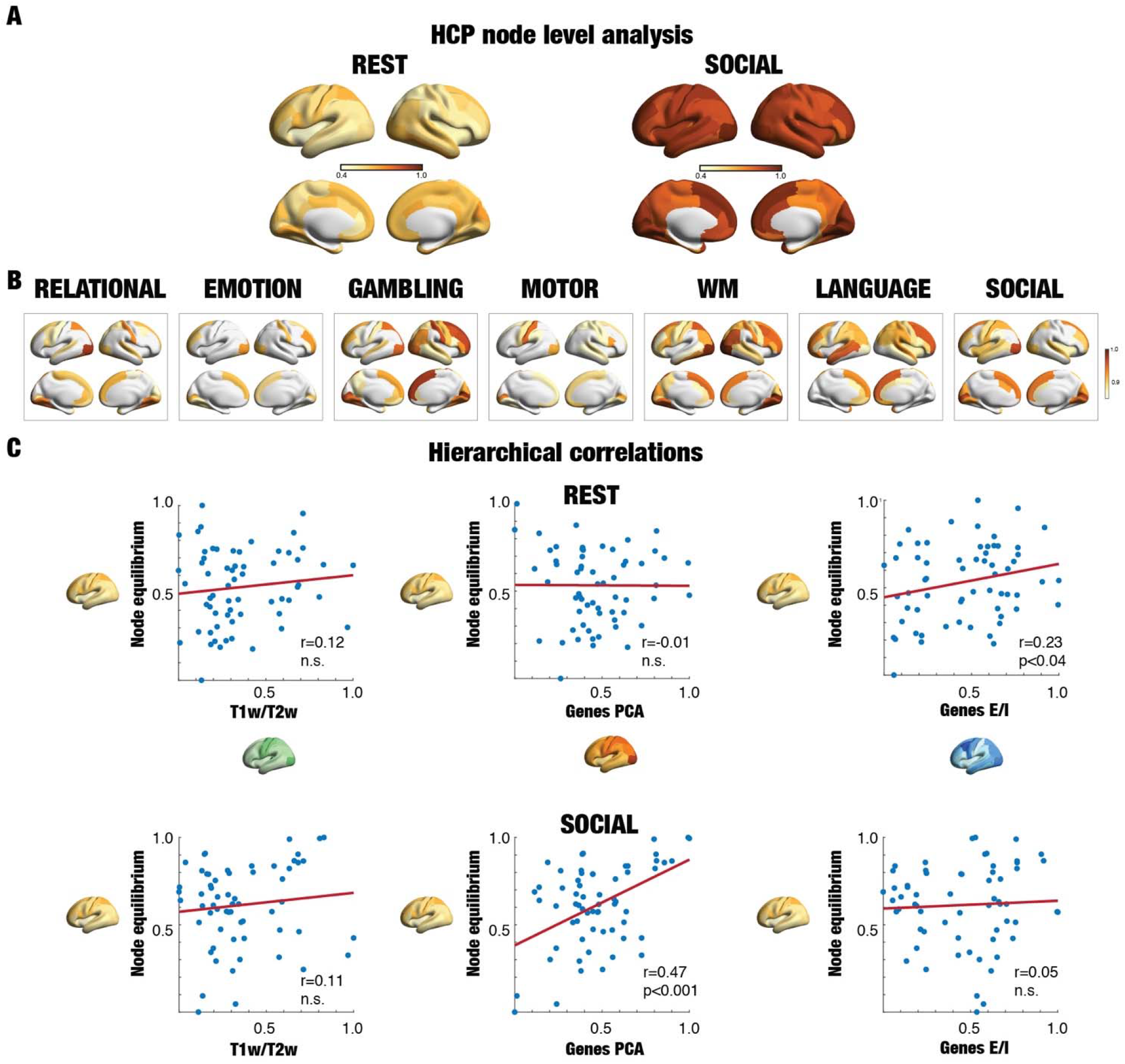
Node-level analysis of HCP data reveals distinct patterns of local non-equilibrium/non-reversibility. Applying TENET at the node-level serves to distinguish the local level of influence on brain regions by the environment. **A)** Rendering of the node-level non-equilibrium for resting and the social task, which show the lowest and highest levels of global non-equilibrium, respectively. This is equally true at the node-level but with significant inter-regional heterogeneity (compare the different shades in a common colormap from yellow to brownish red). **B)** Rendering of the levels of non-equilibrium for all the seven tasks. Notice the clear differences between tasks and how for instance the working memory (WM) task shows high levels of non-equilibrium in prefrontal regions, while the LANGUAGE task shows high levels of non-equilibrium in known language areas consistent with the existing extensive literature. **C)** We studied the hierarchy of the levels of non-equilibrium in rest and tasks by comparing this to other known sources of heterogeneity such as the myelinisation ratio (T1w/T2w ratio, obtained from HCP data) and various forms of gene expressions in the brain, obtained from the Allen Human Brain Atlas (Arnatkeviciute et al., 2019; Deco et al., 2021a; Fornito et al., 2019; Hawrylycz et al., 2012). In particular, we used the first PCA component of all genes and the excitation/inhibition ratio given by the gene-expression for genes coding for the excitatory AMPA and NMDA receptors and inhibitory GABA-A receptor isoforms and subunits. During resting, we found a significantly correlation (p<0.001, non-parametric) with the excitation-inhibition ratio but not with the two others. In contrast, the social task showed a clear significant correlation with the first PCA (p<0.001, non-parametric) but not with the others. These differences between rest and task show how the environment is directly changing the functional hierarchy.

In **Figure 5C** we further investigated these finding demonstrating that our new measure characterises the engagement across the whole brain rather than just in sensory regions. We compared them directly to the myelinisation ratio (T1w/T2w ratio, obtained from HCP data), which contains high values in the sensory regions of visual, somatomotor and auditory (Burt *et al*., 2018). The non-significant correlations between the node levels of non-equilibrium for both REST and SOCIAL task (left row) with this map clearly demonstrates that the new measure is not just linked to sensory but primarily to higher associative brain regions across the whole brain.

Further investigating links to other source of heterogeneity led us to investigate the various forms of gene expressions in the brain as obtained from the Allen Human Brain Atlas (Arnatkeviciute *et al*., 2019; Deco *et al*., 2021a; Fornito *et al*., 2019; Hawrylycz *et al*., 2012). The middle row of **Figure 5C** shows the correlations between and the first PCA component of all genes and the node level of non-equilibrium of REST (top) and SOCIAL task (bottom). Interestingly, there was a significant correlation between the PCA genes values and node levels in the SOCIAL task (r=0.47, p<0.001, non-parametric) but not with the node levels in REST.

We also investigated another major source of heterogeneity, namely the excitation-inhibition (E-I) ratio given by the gene-expression for genes coding for the excitatory AMPA and NMDA receptors and inhibitory GABA-A receptor isoforms and subunits. In contrast to the PCA maps, the rightmost row of **Figure 5C** shows a significant between the node level of non-equilibrium in REST and the E-I values (r=0.23, p<0.04, non-parametric) but not for the node levels in the SOCIAL task.

### Using the arrow of time in neuropsychiatric disease

Given that the TENET framework by design measures how the environment is driving the brain, and its high level of sensitivity demonstrated above, it would appear a promising avenue for better characterising between neuropsychiatric diseases. We therefore applied the TENET framework on the large public UCLA dataset of neuropsychiatric patients with schizophrenia, bipolar and ADHD and matched control group of participants.

**Figure 6** shows the results of using the TENET framework to establish the reversibility on the global and local node levels for the four groups. The left panel of **Figure 6A** shows boxplots of the average reversibility across time, where the control group was significantly higher than each of the neuropsychiatric groups (all p<0.001, Wilcoxon rank sum). This suggests that neuropsychiatric disease reduces the levels of non-equilibrium, suggesting that the brain is less driven by the extrinsic environment. Furthermore, each neuropsychiatric disease group were significantly different from each other (all p<0.001, Wilcoxon rank sum).

**Figure 6.**
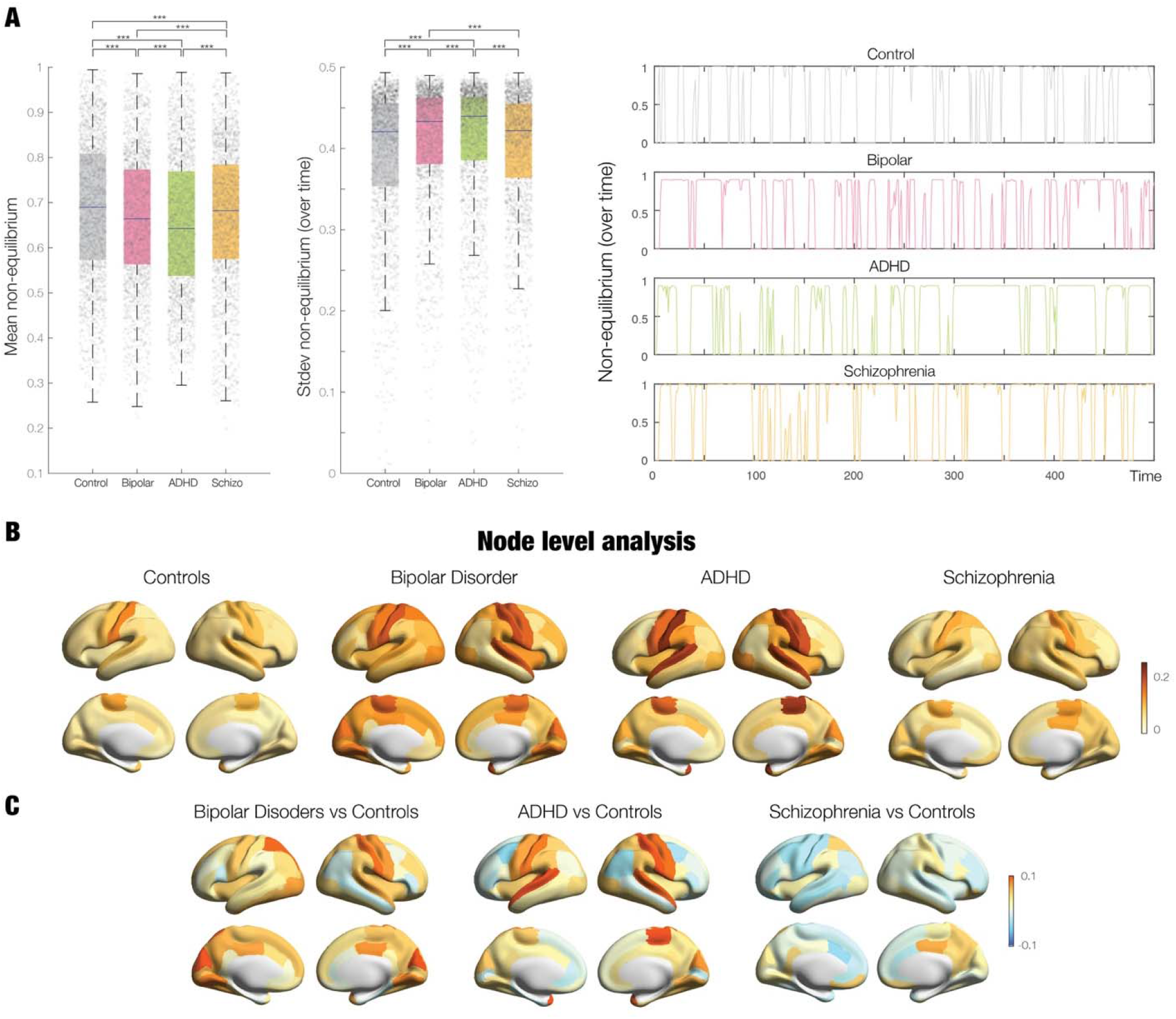
The arrow of time in neuropsychiatric disease. We used the non-equilibrium on the global and local node levels on the large public UCLA dataset of neuropsychiatric patients with schizophrenia, bipolar and ADHD and matched control group of participants. **A)** First, we used the TENET framework to compute the level of non-equilibrium/non-reversibility at the global level for each group. The left panel of boxplots shows that the average reversibility across time for the control group is significantly higher than each of the neuropsychiatric groups (all p<0.001, Wilcoxon rank sum). This suggests that neuropsychiatric disease reduces the levels of non-equilibrium, suggesting that the brain is less driven by the extrinsic environment. In addition, each neuropsychiatric disease group are significantly different from each other (all p<0.001, Wilcoxon rank sum). The middle panel of boxplots shows that the standard deviation of the reversibility across time is significantly reduced for the control participants compared to neuropsychiatric groups and that between them, there are also significant differences (all p<0.001, Wilcoxon rank sum, except for controls vs schizophrenia). The right panel shows examples of the temporal evolution of the reversibility computed by TENET for a participant from each of the four groups. **B)** Complementing these findings at the global level, we used the TENET framework to compute the node level reversibility for each group and show the corresponding thresholded renderings. **C)** In order to stress the differences between the control group and the three neuropsychiatric disorders, we show renderings of these differences. Interestingly, in schizophrenia we found local decreases compared to controls across the brain, primarily located in the temporal, parietal, and prefrontal cortices. In contrast, in bipolar disorder compared to controls, most brain regions except for the somatomotor regions are showing more non-reversibility compared to controls, suggesting that the brain is more driven by the environment. Comparing ADHD with controls shows larger local levels of non-reversibility in somatosensory, temporal, parietal and insular cortices. In particular the somatomotor regions are more driven by the environment, suggesting a route for hyperactivity.

The middle panel of **Figure 6A** shows boxplots of the standard deviation of the reversibility across time for each of the four group, reflecting the levels of non-stationarity. This is significantly reduced for the control participants compared to neuropsychiatric groups and between them (all p<0.001, Wilcoxon rank sum, except controls vs schizophrenia), i.e. the brains of patients with ADHD and bipolar disorder are more non-stationary than controls. To further appreciate the differences between groups, right panel of **Figure 6A** plots examples of the temporal evolution of the global reversibility computed by TENET for all the four groups.

These promising results prompted us to use the TENET framework to compute the mean node level reversibility for each group. **Figure 6B** shows the corresponding renderings on the human brain. We also computed the differences between the mean in the control group with the three neuropsychiatric groups, shown rendered in **Figure 6C**. As can be seen and interpreted in details in the discussion, there are clear differences between groups which suggest that the node level of non-equilibrium might be useful as a biomarker for disease.

## Discussion

We used a thermodynamics-based deep learning Temporal Evolution NETwork (TENET) framework to reveal varying levels of non-reversibility and non-equilibrium in different brain states. At the core of this framework is the ability to quantify how the environment drives the brain by providing a measure of non-equilibrium captured by the arrow of time, i.e. the level of non-reversibility in brain signals. The results of using the TENET framework show that the empirically extracted level of non-reversibility/non-equilibrium can distinguish between different brain states in health and disease.

### Global results

Specifically, we found higher global levels of non-reversibility/non-equilibrium in different tasks than during resting state in the large-scale HCP neuroimaging dataset. Similarly, we showed higher and more variable global levels of non-reversibility/non-equilibrium across time for healthy participants compared to neuropsychiatric patients in the large-scale UCLA neuroimaging dataset.

### Network level results

Complementary to investigating non-equilibrium/non-reversibility at the global level, we aimed to discover if there is more information to be extracted from the network level, specifically regarding differences between rest and task. We used the TENET framework on the seven Yeo resting state networks in the Schaefer 500 parcellation. The results again showed lower levels of non-equilibrium for rest compared to the seven tasks, suggesting that REST is more in equilibrium and therefore more intrinsically driven, while the tasks are clearly driven by the environment as demonstrated by the high levels of non-equilibrium of the sensory networks (VIS and SOM).

Interestingly, the REST condition shows the highest levels of non-equilibrium in the default mode network (DMN) and Visual (VIS) network. Similarly, the sensory networks (VIS and SOM) are showing the highest level of non-equilibrium across the seven tasks (except for LANGUAGE), and thus most driven by the environment. The lowest levels of non-equilibrium are found in the Limbic network (LIM), signalling that the LIM network is more intrinsically driven. Of the seven tasks the lowest levels of non-equilibrium are found in the EMOTION and RELATIONAL tasks, which are almost as low as REST.

The differences across conditions and across networks are quantified *within* conditions in **Figure 4C**, showing significant differences (p<0.05, Wilcoxon rank sum) and *between* conditions in **Figure 4D**. As can be seen most of the comparisons are significant, suggesting that the specificity of level of non-equilibrium/non-reversibility within and between conditions. **Figure 4E** further visualises these differences between conditions by showing a combined spider plot of the levels of non-equilibrium for each Yeo network in all conditions (rest and seven tasks).

Furthermore, using a support vector machine (SVM) with Gaussian kernels to classify the conditions (rest and seven tasks) showed a very high accuracy of 59% compared to the 12.5% chance level. Equally, just classifying rest compared to all tasks the level of accuracy to 93.1%, showing that network-level TENET is an excellent method for distinguishing different cognitive brain states.

### Node-level results

Equally, the TENET framework can be applied at different scales. We investigated this measure at the level of resting state networks and at the node level. Similar to the findings at the global scale, the findings at the local scale reveal clear inter-regional heterogeneities between rest and tasks (see **Figure 5**). The findings show that in many cases higher associative brain regions are more driven by the environment during task performance in the HCP dataset. Specifically, **Figure 5A** shows the node-level rendering for non-equilibrium/non-reversibility for REST and the SOCIAL task. Similar to the results for the global level, these are the two conditions with the lowest and highest levels of non-equilibrium.

The power of the TENET framework is perhaps illustrated in the results for the MOTOR task, where the thresholded results show selective engagement of the somatomotor regions as expected (Barch *et al*., 2013) but equally the results show engagement in visual cortices and midline medial prefrontal regions.

Another good example is the LANGUAGE task, where the results show broad engagement of ventral lateral prefrontal cortex, superior and inferior temporal cortex (including the anterior temporal poles bilaterally (Barch *et al*., 2013; Binder *et al*., 2011). Also, as expected from this primarily auditory task, the visual regions are not in the top 15% of driven regions.

Similarly, the WM task shows high levels of driving in regions including the medial prefrontal cortex, posterior cingulate, and the occipital-parietal junction, fully consistent with the literature (Barch *et al*., 2013; Drobyshevsky *et al*., 2006).

Many previous studies of the HCP dataset have used conventional neuroimaging categorical analyses to bring out the differences between conditions and baseline (Barch *et al*., 2013). Here, we are using the orthogonal approach of the TENET which measures the degree to which the environment drives the brain into non-equilibrium. As a first approximation, our method would be expected to mainly bring out the sensory regions of the brain, given that these regions directly receive external stimulation. Yet, interestingly, the results show that higher associative brain regions are equally and often more driven by the environment during task performance. As such, this could suggest that beyond the level of activation, it is the level to which the environment can drive brain activity as reflected in our measure of non-equilibrium. Even more important, the present framework is not dependent on comparison to a baseline, which has been a main bone of contention in the neuroimaging field since the discovery of intrinsic resting activity (Biswal *et al*., 1995; Biswal *et al*., 2010; Raichle *et al*., 2001).

### Heterogeneity

We further investigated the heterogeneity found at the node-level by comparing the results to various known forms of heterogeneity such as the gene expressions maps obtained from the Allen Human Brain Atlas (Arnatkeviciute *et al*., 2019; Deco *et al*., 2021a; Fornito *et al*., 2019; Hawrylycz *et al*., 2012). We found a significant correlation between the PCA genes values and node levels in the SOCIAL task - but not with the node levels found in REST (see **Figure 5C**). The reverse was true for excitation-inhibition ratio given by the gene-expression, which was correlated with the node levels in REST but not in SOCIAL. These significant differences between rest and task show how the environment is directly changing the functional hierarchy. It is of considerable interest that the E-I ratio is significantly correlated with node levels in REST and not the node levels in task, since this suggest that REST is more intrinsically shaped. On the other hand, the significant correlation between the node levels in the SOCIAL task (but not with REST), suggests that the level of driving by the environment is not fully free but still constrained to a certain degree by genetics (here captured by the first PCA of the genes).

Further investigating the heterogeneity we compared REST and SOCIAL directly to the myelinisation ratio (T1w/T2w ratio), which contains high values in the sensory regions of visual, somatomotor and auditory (Burt *et al*., 2018). Surprisingly, we did not find a correlation to the node levels of non-equilibrium for both REST and SOCIAL task. This result clearly demonstrates that the environment is not driving the brain to non-equilibrium by solely affecting the sensory regions but equally through driving higher associative brain regions across the whole brain. This fits well with the evidence that the three sources of heterogeneity investigated here are correlated among themselves (Burt *et al*., 2018; Deco *et al*., 2021a), yet influencing the functional hierarchy at the node-level of non-equilibrium differently under different conditions. Overall, these results provide evidence for the severe constraints in terms of degrees of freedom that the brain has to operate within.

### UCLA node level

In addition to the lower global levels of non-equilibrium/non-reversibility found in patients with neuropsychiatric disease compared to healthy participants, we also found significant local heterogenous node-level changes differentiating between the different disorders. Interestingly, in schizophrenia compared to controls, we found local decreases across the brain but primarily located in the temporal, parietal, and prefrontal cortices. These regions are clearly less driven by the environment, which is compatible with the literature showing that the disorder is associated with more isolation, as a function of the loss of balance between intrinsic and extrinsic activity (van den Heuvel and Fornito, 2014).

In contrast, in bipolar disorder compared to controls most brain regions except for the somatomotor regions are showing more non-reversibility compared to controls, suggesting that the brain is more driven by the environment, which is consistent with the literature suggesting the large sudden swings in mood (Furman *et al*., 2011; Menon and Uddin, 2010). Comparing ADHD with controls shows larger local levels of non-reversibility in somatosensory, temporal, parietal and insular cortices. In particular the somatomotor regions are more driven by the environment, suggesting a route for hyperactivity, while the lower non-reversibility in parietal regions could be linked to the known attentional deficits in the disorder (Konrad and Eickhoff, 2010).

### TENET components

The TENET framework relies on two essential elements; namely the concept of reversibility (as captured by the arrow of time) and how deep learning is able to quantify the reversibility of brain signals. In terms of the arrow of time, this popularisation of this idea is usually credited to the physicist Arthur Eddington (Eddington, 1928). Here we showed that this key idea from physics and thermodynamics can equally well be applied in neuroscience. The second law of thermodynamics, as immortalised by Rudolph Clausius (Clausius, 1865) and Sadi Carnot (Carnot, 1824) states a non-equilibrium is characterised by the arrow of time which indicates the non-reversibility of a system. In fact, the second law of thermodynamics can be expressed by the Clausius inequality which establishes that the work associated with the process (averaged over many repetitions) is larger than the change in its free energy, which is the same as stating that the system is non-reversible and in non-equilibrium.

In terms of deep learning, this method has received a lot of attention over the last couple of years. This powerful machine learning technique has proven highly useful for providing solutions to a number of difficult computational problems ranging from vision to playing Go (Sejnowski, 2018, 2020; Silver *et al*., 2018; Yang and Wang, 2020). However, some criticisms have been raised over the largely black-box nature of these advances, which have had considerable practical utility for solving complex problems, but have produced little in way of new insight into *how* this is achieved mechanistically (Marcus, 2018). Recent research has, however, started to harness the power of deep learning for discovering useful underlying mechanisms (Seif *et al*., 2021).

Here, we were not aiming to use deep learning as a technique for revealing underlying brain mechanisms but rather simply as a tool for providing the level of the reversibility of the arrow of time in brain signals. In other words, in our framework the important question is to determine the level of distinction between forward and time-reversed back time series, but not *how* this is achieved. As such, here the black box nature of deep learning was not relevant for solving the non-trivial problem of determining non-reversibility/non-equilibrium.

As mentioned above, the TENET framework can be studied at all spatial scales. Here, however, we focused on three levels: global, network and node levels, where the global level considers all the whole-brain signals, while the node level considers the signals in each brain region separately and the network level considers the typical large-scale resting state networks (Biswal *et al*., 1995; Biswal *et al*., 2010; Smith *et al*., 2013a). This allowed us to focus on different aspects of non-equilibrium, i.e. the interactions between the environment and different brain scales, ranging from whole-brain to large-scale networks and to regions. Indeed, the node level turned out to be a highly sensitive measure of quantifying and interpreting different cognitive brain states and differences between health and multiple diseases.

### Perspectives

This thermodynamics-based deep learning framework opens up for an examination of differences of non-reversibility/non-equilibrium for different levels of consciousness; from sleep and anaesthesia to altered states of consciousness induced by psychedelics and meditation. The framework can easily be extended to other neuroimaging modalities such as MEG/EEG. It can even be used with LFP and other types of cell recordings in animals.

One particularly challenging question relates to the role of brain processing of reversible external stimuli such as for example watching the forward and backward versions of the movie of a glass being shattered. How will the non-reversibility/non-equilibrium in brain dynamics change with identical stimuli but where the order has been changed such that the arrow of time has been violated? Will this elicit different non-reversibility/non-equilibrium in brain dynamics when showing the forward and backward versions of a movie of billiards balls moving, which is not in any clear way violating the arrow of time? These stimuli are experienced radically different, which must be linked to the interactions between the extrinsic stimulation with the dynamics of intrinsic predictability formed by prior experiences. However, intrinsic predictability is known to be affected in neuropsychiatric disease and it would be of considerable interest to study how the reversibility of external stimulation influences the non-reversibility/non-equilibrium in brain dynamics. In fact, this could potentially reveal new information about the interactions between intrinsic and extrinsic dynamics which are known to be compromised. The reversal of external stimuli could also take place at higher cognitive levels, such as when inverting the nodes of Bach’s fugue or that of a complex narrative.

From its conception, the arrow of time is coupled to the deep notion of causality. Thermodynamics offers important tools for establishing the causal directionality of information flow through the concept reversibility and entropy. There is of course a large literature on causality, best summarised in the seminal work by Judea Pearl (Pearl, 2009), where he shows that any framework of causal inference is based on inferring causal structures that are equivalent in terms of the probability distributions they generate; that is, they are indistinguishable from observational data, and could only be distinguished by manipulating the whole system.

In neuroscience, there has been numerous attempts to capture causality in brain dynamics. One influential concept is ignition, the idea that a stimulus can ignite a causal chain of events propagating across the brain (1998). This ignition can happen as a result of extrinsic stimulation (Joglekar *et al*., 2018; Mashour *et al*., 2020; van Vugt *et al*., 2018) or as part of intrinsic events (Deco *et al*., 2021a; Deco *et al*., 2017). More sophisticated approaches uses probabilistic principles of mutual information (Brovelli *et al*., 2015; Pereda *et al*., 2005; Quiroga *et al*., 2000; Quiroga *et al*., 2002) to determine the directional causality underlying the functional hierarchical organisation of brain function (Deco *et al*., 2021b).

The concept of the arrow of time has also been investigated from the perspective of chaos theory, originating with the work of Henri Poincaré who published the first description of chaotic motion in 1890 (Poincaré, 1890). Later work has confirmed that one key characteristic of chaos is the infinite sensitivity to initial conditions (Strogatz, 2018). Given this extreme sensitivity, even if a classic mechanic deterministic chaotic system is in principle reversible, in practice this is in fact non-reversible. In other words, chaos makes it very difficult to establish the computational reversibility and thus causality. Turbulence is a classic example of a spatiotemporal chaotic system which is associated with non-equilibrium and thus non-reversibility. Interestingly, turbulence is a highly useful dynamical regime for optimally transferring energy/information over spacetime and it has recently been shown that brain dynamics are indeed turbulent (Deco and Kringelbach, 2020). The turbulent regime supports the information cascade which is crucial for extracting order from disorder.

Overall, the novel thermodynamics-based deep learning TENET framework can provide detailed information of the varying levels of non-stationary and non-equilibrium nature of brain dynamics in health and disease driven by the environment. The TENET framework offers a quantitative account of differences in non-reversibility/non-equilibrium. Future work could integrate this with causal mechanistic whole-brain modelling in a turbulent regime to deepen our understanding of how brain dynamics organise human behaviour in the face of the second law of thermodynamics in health and disease.

## Methods

### Neuroimaging Ethics

For the HCP dataset, the Washington University–University of Minnesota (WU-Minn HCP) Consortium obtained full informed consent from all participants, and research procedures and ethical guidelines were followed in accordance with Washington University institutional review board approval.

For the UCLA dataset, as detailed in (Poldrack *et al*., 2016), the Consortium for Neuropsychiatric Phenomics recruited neuropsychiatric participants and healthy controls who gave written informed consent following procedures approved by the Institutional Review Boards at UCLA and the Los Angeles County Department of Mental Health.

### Neuroimaging Participants HCP rest and task

The data set used for this investigation was selected from the March 2017 public data release from the Human Connectome Project (HCP) where we chose a sample of 990 participants from the total of 1003 participants, since not all participants performed all tasks.

### The HCP task battery of seven tasks

The HCP task battery consists of seven tasks: working memory, motor, gambling, language, social, emotional, relational, which are described in details on the HCP website (Barch *et al*., 2013). HCP states that the tasks were designed to cover a broad range of human cognitive abilities in seven major domains that sample the diversity of neural systems 1) visual, motion, somatosensory, and motor systems, 2) working memory, decision-making and cognitive control systems; 3) category-specific representations; 4) language processing; 5) relational processing; 6) social cognition; and 7) emotion processing. In addition to resting state scans, all 1003 HCP participants performed all tasks in two separate sessions (first session: working memory, gambling and motor; second session: language, social cognition, relational processing and emotion processing). As a test-retest control condition, a small subsample of 45 HCP participants performed the paradigm twice.

### Neuroimaging Participants UCLA rest

Consortium for Neuropsychiatric Phenomics published a dataset with neuroimaging as well as phenotypic information for 272 participants. We used the preprocessed data with a total of 261 participants, since seven of the participants were missing T1 weighted scans (Gorgolewski *et al*., 2017) and three healthy controls and one ADHD patient were missing resting state scans. The total population analysed consists of 122 healthy controls, as well as participants with diagnoses of adult ADHD (40 patients), bipolar disorder (49 patients) and schizophrenia (50 patients).

### Neuroimaging structural connectivity and extraction of functional timeseries

#### HCP preprocessing and extraction of functional timeseries in fMRI resting state and task data

The preprocessing of the HCP resting state and task datasets is described in details on the HCP website. Briefly, the data is preprocessed using the HCP pipeline which is using standardized methods using FSL (FMRIB Software Library), FreeSurfer, and the Connectome Workbench software (Glasser *et al*., 2013; Smith *et al*., 2013b). This standard preprocessing included correction for spatial and gradient distortions and head motion, intensity normalization and bias field removal, registration to the T1 weighted structural image, transformation to the 2mm Montreal Neurological Institute (MNI) space, and using the FIX artefact removal procedure (Navarro Schroder *et al*., 2015; Smith *et al*., 2013b). The head motion parameters were regressed out and structured artefacts were removed by ICA+FIX processing (Independent Component Analysis followed by FMRIB’s ICA-based X-noiseifier (Griffanti *et al*., 2014; Salimi-Khorshidi *et al*., 2014)). Preprocessed timeseries of all grayordinates are in HCP CIFTI grayordinates standard space and available in the surface-based CIFTI file for each participants for resting state and each of the seven tasks.

We used a custom-made Matlab script using the ft_read_cifti function (Fieldtrip toolbox (Oostenveld *et al*., 2011)) to extract the average timeseries of all the grayordinates in each region of the DK80 parcellation, which are defined in the HCP CIFTI grayordinates standard space. The BOLD timeseries were filtered using a second-order Butterworth filter in the range of 0.008-0.08Hz.

#### UCLA preprocessing and extraction of functional timeseries in fMRI resting state data

The preprocessing of UCLA resting state datasets is described in details on the website and in the paper by Gorgolewski and colleagues (Gorgolewski *et al*., 2017). Briefly, the preprocessing was performed using FMRIPREP version 0.4.4 (http://fmriprep.readthedocs.io). This robust preprocessing pipeline is based on the Nipype workflow engine7 and aims to combine different implementations of various MR signal processing algorithms (from established software packages) to deliver a robust spatial normalization and nuisance estimation workflow.

#### Parcellations

For the analysis at the global and node-level, all neuroimaging data was processed using the DK80 standard parcellations (Deco *et al*., 2021b). Briefly, this was constructed using the Mindboggle-modified Desikan-Killiany parcellation (Desikan *et al*., 2006) with a total of 62 cortical regions (31 regions per hemisphere) (Klein and Tourville, 2012). We added the 18 subcortical regions, ie nine regions per hemisphere: hippocampus, amygdala, subthalamic nucleus (STN), globus pallidus internal segment (GPi), globus pallidus external segment (GPe), putamen, caudate, nucleus accumbens and thalamus. This provided a total of 80 regions in the DK80 parcellation; also precisely defined in the common HCP CIFTI grayordinates standard space. For the analysis at the network level we also used the Schaefer500 where each parcel is marked with the seven resting state network in the Yeo7 parcellation (Schaefer *et al*., 2018; Yeo *et al*., 2011).

### TENET deep learning framework and associated methods

The TENET deep learning framework is a general method which can use many types of data. However, it is important that the data for training and generalisation are balanced between different conditions to We used TENET to carry out two independent analyses using two different datasets from HCP and UCLA and below we show how we avoided any potential problems with bias.

#### HCP dataset

For the large-scale HCP dataset, we performed the reversibility and non-equilibrium analyses at three different spatial scales: *Global, network* and *local node*-*level*. For each scale, we trained and analysed three different TENETs, one for each spatial scale. We used roughly 90% of the data for the training set (890 participants) and the remaining 10% for the test and validation sets (100 participants).

For the global level, the input patterns consisted of the windowed timeseries across the whole brain in the DK80 parcellation, i.e. each pattern consisted of 80 windowed timeseries.

For the network level, the input patterns consisted of the windowed timeseries in the Schaefer500 parcellation, i.e. which each parcel belong to one of the seven Yeo resting state networks. We performed the analysis for the Schaefer500 parcels belonging to each of the seven Yeo networks and thus obtained a measure of non-equilibrium for each Yeo network.

Finally, for the finest spatial scale at the node level, the input patterns consisted of the windowed timeseries, for one parcel in the DK80 parcellation, i.e. each of the patterns consisted of one windowed timeseries for that parcel. We performed this for each of the 80 parcels in the DK80 separately such that we obtained a measure of non-equilibrium for each brain region. For the analysis of the HCP data, we carried out these three spatial scale analyses for HCP rest and seven tasks, i.e. a total of 24 different analyses.

The input to TENET consisted of sliding windows from the BOLD neuroimaging data from all the participants, shortened to the shortest duration of a task (EMOTION task with 176 TRs). We generated these sliding windows with slightly different parameters for the three types of analysis: For global and network level analysis, we used a windows size of 20 TRs (each lasting 0.72 seconds) and for node-level analysis, we used a window size of 150 TRs. These were then shifted forward until the end of the datasets (with a length of 3 TRs for global and network level analysis and 2 TRs for node-level analysis).

For each of the sliding windows, we generated two input patterns containing 1) forward and 2) time reversed backward sliding windowed timeseries and for the supervised learning, we associated each pattern with the output class label of forward and backward, respectively.

#### UCLA dataset

For the analysis of the smaller scale UCLA dataset with coarser TR (2 seconds) and shorter duration (152 TRs), we carried out different spatial scale analyses at the level of global and node-level for the resting state data of control and three neuropsychiatric disorders (schizophrenia, bipolar disorder and ADHD). For this much smaller dataset of 261 participants, we used roughly 90% of the data for the training set (40 participants in each of the four conditions) and the remaining data for the test and validation sets (i.e. 36 participants for the global and 35 participants for node-level analysis). For the global analysis, we shuffled the data 500 times in order to perform non-parametric significance testing (for the boxplot in **Figure 6A**). For the node analysis we shuffled the data 10 times and computed the average node-level of non-equilibrium/non-reversibility.

The input to TENET consisted of sliding windows from the BOLD neuroimaging data from all the participants. We generated these sliding windows with slightly different parameters for the two types of analysis: For global level analysis, we used a windows size of 10 TRs of 2 seconds and for node-level analysis, we used a window size of 20 TRs. These were then shifted forward until the end of the dataset with a length of 1 TR.

#### Training and generalisation in deep learning network TENET

This data was then used with TENET for training and generalisation in the following way. After training, we measure the reversibility of each pattern in the validation set. This is computed by comparing the trained output for the forward and backward versions of each pattern. More specifically, for all three spatial scales, we compute the level of reversibility, *R*(*t*), for a given sliding window at time t by using the following equation

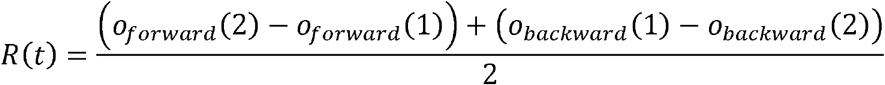

Here *o*_*forward*_(*i*) is the output *i* of the final output classification layer, when the forward pattern is presented in the input layer. Similarly, *o*_*backward*_(*i*) is the output for node *i* of the final output classification layer but now when a backward pattern is presented in the input layer. Given that the forward and backward categories are associated with the outputs *o*_*forward*_ = [0,1] and *o*_*backward*_ = [1,0], respectively, *R*(*t*) thus represents the degree of reversibility for a particular sliding window at time *t*. If either the degrees of right classification of forward (*o*_*forward*_(2) − *o*_*forward*_(1)) or backward (*o*_*backward*_(1) − o_*backward*_(2)) are smaller than 0, this means that the classification is incorrect and in this case we set *R*(*t*) = 0. The degree of reversibility *R*(*t*) is measuring the degree of non-equilibrium at time *t*. We also report the mean and the standard deviation over time, which corresponds to what is called the mean non-equilibrium and the standard deviation of non-equilibrium (over time) in the results and figures.

For the architecture of TENET, we use the standard MATLAB architecture with an input layer of size *N* * *w*, where *N* corresponds to the spatial scale (i.e. global *N* = 80, network *N* = 7, node level *N* = 1) and *w* is the size of the sliding window. This is followed by 10 fully connected layers including 1) batch normalisation, which normalises a mini-batch of data across all observations for each channel independently and 2) a non-linearity operation implemented using *reluLayer*, which performs a threshold operation to each element of the input, where any value less than zero is set to zero. The dimensions of the 10 layers are [2048;1024;512;256;128;64;32;16;8;4]. The final layer of TENET is a softmax classification layer of dimension [2], corresponding the two possible output class labels (forward and backward).

For training TENET, we used the deep learning algorithm ADAM, which is an algorithm for first-order gradient-based optimization of stochastic objective functions, based on adaptive estimates of lower-order moments to attenuate the effects of noise as it is implemented in MATLAB with the recommended default parameters (Kingma and Ba, 2014).

#### Measuring the breaking of detailed balance of the system

We wanted to test if non-equilibrium is associated with the breaking of detailed balance of a system. In order to show the increase in non-balance in non-equilibrium, we decided to characterise the level of asymmetry in the causal interaction as expressed by the shifted correlation matrix. Specifically, we selected patterns from sliding windows of low and high reversibility, and computed the time-delayed functional connectivity matrix between all pairs of brain regions, *FC*(τ)

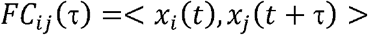

Where < > indicates correlation over time and we used τ =3 (in TRs). We computed the *FC*_*ij*_(τ) over all participants and all sliding windows for each condition (for HCP REST and the SOCIAL task, the most non-equilibrium task). For the global level of asymmetry, we computed for each sliding window the degree of asymmetry as the mean value of the elements of the matrix: *FC*_*ij*_(τ) − *FC*_*ij*_ (τ)^*T*^, where the superscript T indicates the transposition. For the node-level of asymmetry, we first computed for each sliding window, the incoming flow *F*_*in*_(*i*) = ∑_*j*_ *FC*_*ij*_ (τ), and the outcoming flow *F*_*out*_(*j*) = ∑_*i*_*FC*_*ij*_ (τ). The node-level of asymmetry for node *i* is then given by < |*F*_*out*_(*i*) − *F*_*in*_(*i*)| >_*t*_, where < >_*t*_ indicates the average over all sliding windows and participants. We render the change between high and low levels of the node-level asymmetry.

#### Support vector machine for network level classification

We used a support vector machine (SVM) with Gaussian kernels as implemented in the Matlab function *fitcecoc*. The function returns a full, trained, multiclass, error-correcting output codes (ECOC) model. This is achieved using the predictors in the input with class labels. The function uses K(K – 1)/2 binary SVM models using the one-versus-one coding design, where we used K=8 as the number of unique class labels. In other words, the SVM had seven inputs (the Yeo resting state networks) corresponding to the output produced by the network level TENET. The output was eight classes corresponding to the conditions (rest and seven tasks). We used the output from the 100 HCP participants used for generalisation, subdivided into 90% training and 10% validation, repeated and shuffled 100 times.

## Data availability

The multimodal neuroimaging data are freely available from HCP. The UCLA data are freely available from the Consortium for Neuropsychiatric Phenomics.

## Code availability

The code used to run the analysis is available on GitHub (https://github.com/decolab/tenet).

## Acknowledgements

G.D. is supported Spanish national research project (ref. PID2019-105772GB-I00 MCIU AEI) funded by the Spanish Ministry of Science, Innovation and Universities (MCIU), State Research Agency (AEI); HBP SGA3 Human Brain Project Specific Grant Agreement 3 (grant agreement no. 945539), funded by the EU H2020 FET Flagship programme; SGR Research Support Group support (ref. 2017 SGR 1545), funded by the Catalan Agency for Management of University and Research Grants (AGAUR); Neurotwin Digital twins for model-driven non-invasive electrical brain stimulation (grant agreement ID: 101017716) funded by the EU H2020 FET Proactive programme; euSNN European School of Network Neuroscience (grant agreement ID: 860563) funded by the EU H2020 MSCA-ITN Innovative Training Networks; CECH The Emerging Human Brain Cluster (Id. 001-P-001682) within the framework of the European Research Development Fund Operational Program of Catalonia 2014-2020; Brain-Connects: Brain Connectivity during Stroke Recovery and Rehabilitation (id. 201725.33) funded by the Fundacio La Marato TV3; Corticity, FLAG–ERA JTC 2017 (ref. PCI2018-092891) funded by the Spanish Ministry of Science, Innovation and Universities (MCIU), State Research Agency (AEI). MLK is supported by the Center for Music in the Brain, funded by the Danish National Research Foundation (DNRF117), and Centre for Eudaimonia and Human Flourishing funded by the Pettit and Carlsberg Foundations.

## References

Arnatkeviciute, A., Fulcher, B. D. and Fornito, A. (2019) A practical guide to linking brain-wide gene expression and neuroimaging data. NeuroImage 189, 353–367.

Attwell, D. and Laughlin, S. B. (2001) An energy budget for signaling in the grey matter of the brain. Journal of Cerebral Blood Flow & Metabolism 21, 1133–1145.

Barch, D. M., Burgess, G. C., Harms, M. P., Petersen, S. E., Schlaggar, B. L., Corbetta, M., Glasser, M. F., Curtiss, S., Dixit, S., Feldt, C., Nolan, D., Bryant, E., Hartley, T., Footer, O., Bjork, J. M., Poldrack, R., Smith, S., Johansen-Berg, H., Snyder, A. Z., Van Essen, D. C. and Consortium, W. U.-M. H. (2013) Function in the human connectome: task-fMRI and individual differences in behavior. NeuroImage 80, 169–189.

Binder, J. R., Gross, W. L., Allendorfer, J. B., Bonilha, L., Chapin, J., Edwards, J. C., Grabowski, T. J., Langfitt, J. T., Loring, D. W., Lowe, M. J., Koenig, K., Morgan, P. S., Ojemann, J. G., Rorden, C., Szaflarski, J. P., Tivarus, M. E. and Weaver, K. E. (2011) Mapping anterior temporal lobe language areas with fMRI: a multicenter normative study. NeuroImage 54, 1465–1475.

Biswal, B., Yetkin, F., Haughton, V. and Hyde, J. (1995) Functional connectivity in the motor cortex of resting human brain using echo-planar MRI. Magnetic resonance in medicine : official journal of the Society of Magnetic Resonance in Medicine / Society of Magnetic Resonance in Medicine 34, 537–541.

Biswal, B. B., Mennes, M., Zuo, X. N., Gohel, S., Kelly, C., Smith, S. M., Beckmann, C. F., Adelstein, J. S., Buckner, R. L., Colcombe, S., Dogonowski, A. M., Ernst, M., Fair, D., Hampson, M., Hoptman, M. J., Hyde, J. S., Kiviniemi, V. J., Kotter, R., Li, S. J., Lin, C. P., Lowe, M. J., Mackay, C., Madden, D. J., Madsen, K. H., Margulies, D. S., Mayberg, H. S., McMahon, K., Monk, C. S., Mostofsky, S. H., Nagel, B. J., Pekar, J. J., Peltier, S. J., Petersen, S. E., Riedl, V., Rombouts, S. A., Rypma, B., Schlaggar, B. L., Schmidt, S., Seidler, R. D., Siegle, G. J., Sorg, C., Teng, G. J., Veijola, J., Villringer, A., Walter, M., Wang, L., Weng, X. C., Whitfield-Gabrieli, S., Williamson, P., Windischberger, C., Zang, Y. F., Zhang, H. Y., Castellanos, F. X. and Milham, M. P. (2010) Toward discovery science of human brain function. Proceedings of the National Academy of Sciences of the United States of America 107, 4734–4739.

Brovelli, A., Chicharro, D., Badier, J. M., Wang, H. and Jirsa, V. (2015) Characterization of Cortical Networks and Corticocortical Functional Connectivity Mediating Arbitrary Visuomotor Mapping. The Journal of neuroscience : the official journal of the Society for Neuroscience 35, 12643–12658.

Burt, J. B., Demirtas, M., Eckner, W. J., Navejar, N. M., Ji, J. L., Martin, W. J., Bernacchia, A., Anticevic, A. and Murray, J. D. (2018) Hierarchy of transcriptomic specialization across human cortex captured by structural neuroimaging topography. Nature neuroscience 21, 1251–1259.

Carnot, S. (1824) Reflections on the motive power of fire, and on machines fitted to develop that power. Paris: Bachelier 108.

Clarke, D. D. and Sokoloff, L. (1999) Circulation and energy metabolism of the brain. In: Basic Neurochemistry. Molecular, Cellular and Medical Aspects (6th edn). pp. 637–670. Eds. B. W. Agranoff, G. J. Siegel. Lippincott-Raven.

Clausius, R. (1865) Ueber verschiedene für die Anwendung bequeme Formen der Hauptgleichungen der mechanischen Wärmetheorie (Vorgetragen in der naturforsch. Gesellschaft zu Zürich den 24. April 1865). Annalen der Physik und Chemie 125, 353–400.

Collin, D., Ritort, F., Jarzynski, C., Smith, S. B., Tinoco, I. and Bustamante, C. (2005) Verification of the Crooks fluctuation theorem and recovery of RNA folding free energies. Nature 437, 231–234.

Crooks, G. E. (1998) Nonequilibrium measurements of free energy differences for microscopically reversible Markovian systems. Journal of Statistical Physics 90, 1481–1487.

Deco, G., Kringelbach, M., Arnatkevičiūtė, A., Oldham, S., Sabaroedin, K., Rogasch, N. C., Aquino, K. and Fornito, A. (2021a) Dynamical consequences of regional heterogeneity in the brain’s transcriptional landscape. Science Advances, in press.

Deco, G. and Kringelbach, M. L. (2020) Turbulent-like dynamics in the human brain. Cell Reports 33, 108471.

Deco, G., Tagliazucchi, E., Laufs, H., Sanjuán, A. and Kringelbach, M. L. (2017) Novel intrinsic ignition method measuring local-global integration characterises wakefulness and deep sleep. eNeuro 4, e0106-0117.2017.

Deco, G., Vidaurre, D. and Kringelbach, M. L. (2021b) Revisiting the Global Workspace orchestrating the hierarchical organisation of the human brain. Nature Human Behaviour.

Dehaene, S., Kerszberg, M. and Changeux, J. P. (1998) A neuronal model of a global workspace in effortful cognitive tasks. Proceedings of the National Academy of Sciences of the United States of America 95, 14529–14534.

Desikan, R. S., Se, F., Fischl, B., Quinn, B. T., Dickerson, B. C., Blacker, D., Buckner, R. L., Dale, A. M., Maguire, R. P., Hyman, B. T., Albert, M. S. and Killiany, R. J. (2006) An automated labeling system for subdividing the human cerebral cortex on MRI scans into gyral based regions of interest. NeuroImage 31, 968–980.

Drobyshevsky, A., Baumann, S. B. and Schneider, W. (2006) A rapid fMRI task battery for mapping of visual, motor, cognitive, and emotional function. NeuroImage 31, 732–744.

Eddington, A. S. (1928) The Nature of the Physical World. Macmillan: London.

Feng, E. H. and Crooks, G. E. (2008) Length of time’s arrow. Physical review letters 101, 090602.

Fornito, A., Arnatkeviciute, A. and Fulcher, B. D. (2019) Bridging the Gap between Connectome and Transcriptome. Trends in cognitive sciences 23, 34–50.

Frisch, U. (1995) Turbulence: The Legacy of A. N. Kolmogorov. Cambridge University Press: Cambridge.

Furman, D. J., Hamilton, J. P. and Gotlib, I. H. (2011) Frontostriatal functional connectivity in major depressive disorder. Biology of Mood & Anxiety Disorders 1, 11.

Glasser, M. F., Sotiropoulos, S. N., Wilson, J. A., Coalson, T. S., Fischl, B., Andersson, J. L., Xu, J., Jbabdi, S., Webster, M., Polimeni, J. R., Van Essen, D. C., Jenkinson, M. and Consortium, W. U.-M. H. (2013) The minimal preprocessing pipelines for the Human Connectome Project. NeuroImage 80, 105–124.

Gnesotto, F. S., Mura, F., Gladrow, J. and Broedersz, C. P. (2018) Broken detailed balance and non-equilibrium dynamics in living systems: a review. Reports on Progress in Physics 81, 066601.

Gorgolewski, K. J., Durnez, J. and Poldrack, R. A. (2017) Preprocessed Consortium for Neuropsychiatric Phenomics dataset. F1000Res 6, 1262.

Griffanti, L., Salimi-Khorshidi, G., Beckmann, C. F., Auerbach, E. J., Douaud, G., Sexton, C. E., Zsoldos, E., Ebmeier, K. P., Filippini, N., Mackay, C. E., Moeller, S., Xu, J., Yacoub, E., Baselli, G., Ugurbil, K., Miller, K. L. and Smith, S. M. (2014) ICA-based artefact removal and accelerated fMRI acquisition for improved resting state network imaging. NeuroImage 95, 232–247.

Hawrylycz, M. J., Lein, E. S., Guillozet-Bongaarts, A. L., Shen, E. H., Ng, L., Miller, J. A., van de Lagemaat, L. N., Smith, K. A., Ebbert, A., Riley, Z. L., Abajian, C., Beckmann, C. F., Bernard, A., Bertagnolli, D., Boe, A. F., Cartagena, P. M., Chakravarty, M. M., Chapin, M., Chong, J., Dalley, R. A., David Daly, B., Dang, C., Datta, S., Dee, N., Dolbeare, T. A., Faber, V., Feng, D., Fowler, D. R., Goldy, J., Gregor, B. W., Haradon, Z., Haynor, D. R., Hohmann, J. G., Horvath, S., Howard, R. E., Jeromin, A., Jochim, J. M., Kinnunen, M., Lau, C., Lazarz, E. T., Lee, C., Lemon, T. A., Li, L., Li, Y., Morris, J. A., Overly, C. C., Parker, P. D., Parry, S. E., Reding, M., Royall, J. J., Schulkin, J., Sequeira, P. A., Slaughterbeck, C. R., Smith, S. C., Sodt, A. J., Sunkin, S. M., Swanson, B. E., Vawter, M. P., Williams, D., Wohnoutka, P., Zielke, H. R., Geschwind, D. H., Hof, P. R., Smith, S. M., Koch, C., Grant, S. G. N. and Jones, A. R. (2012) An anatomically comprehensive atlas of the adult human brain transcriptome. Nature 489, 391–399.

Jarzynski, C. (2011) Equalities and inequalities: Irreversibility and the second law of thermodynamics at the nanoscale. Annu. Rev. Condens. Matter Phys. 2, 329–351.

Joglekar, M. R., Mejias, J. F., Yang, G. R. and Wang, X. J. (2018) Inter-areal Balanced Amplification Enhances Signal Propagation in a Large-Scale Circuit Model of the Primate Cortex. Neuron 98, 222–234 e228.

Kingma, D. P. and Ba, J. (2014) Adam: A method for stochastic optimization. arXiv preprint arXiv:1412.6980.

Klein, A. and Tourville, J. (2012) 101 labeled brain images and a consistent human cortical labeling protocol. Frontiers in neuroscience 6, 171.

Konrad, K. and Eickhoff, S. B. (2010) Is the ADHD brain wired differently? A review on structural and functional connectivity in attention deficit hyperactivity disorder. Human brain mapping 31, 904–916.

Lynn, C. W., Cornblath, E. J., Papadopoulos, L., Bertolero, M. A. and Bassett, D. S. (2020) Broken detailed balance and entropy production in the human brain. arXiv preprint arXiv:2005.02526.

Magistretti, P. J., Pellerin, L., Rothman, D. L. and Shulman, R. G. (1999) Energy on demand. Science 283, 496–497.

Maragakis, P., Ritort, F., Bustamante, C., Karplus, M. and Crooks, G. E. (2008) Bayesian estimates of free energies from nonequilibrium work data in the presence of instrument noise. The Journal of chemical physics 129, 07B609.

Marcus, G. (2018) Deep learning: A critical appraisal. arXiv preprint arXiv:1801.00631.

Mashour, G. A., Roelfsema, P., Changeux, J.-P. and Dehaene, S. (2020) Conscious processing and the global neuronal workspace hypothesis. Neuron 105, 776–798.

Menon, V. and Uddin, L. Q. (2010) Saliency, switching, attention and control: A network model of insula function. Brain structure & function 214, 655–667.

Navarro Schroder, T., Haak, K. V., Zaragoza Jimenez, N. I., Beckmann, C. F. and Doeller, C. F. (2015) Functional topography of the human entorhinal cortex. eLife 4.

Oostenveld, R., Fries, P., Maris, E. and Schoffelen, J. M. (2011) FieldTrip: Open source software for advanced analysis of MEG, EEG, and invasive electrophysiological data. Comput Intell Neurosci 2011, 156869.

Palus, M. (1996) Nonlinearity in normal human EEG: cycles, temporal asymmetry, nonstationarity and randomness, not chaos. Biological cybernetics 75, 389–396.

Pearl, J. (2009) Causality. Models, reasoning and inference. Cambridge University Press: Cambridge.

Pereda, E., Quiroga, R. Q. and Bhattacharya, J. (2005) Nonlinear multivariate analysis of neurophysiological signals. Prog Neurobiol 77, 1–37.

Poincaré, H. (1890) Sur le problème des trois corps et les équations de la dynamique. Acta mathematica 13, A3–A270.

Poldrack, R. A., Congdon, E., Triplett, W., Gorgolewski, K. J., Karlsgodt, K. H., Mumford, J. A., Sabb, F. W., Freimer, N. B., London, E. D., Cannon, T. D. and Bilder, R. M. (2016) A phenome-wide examination of neural and cognitive function. Scientific data 3, 160110.

Quiroga, R. Q., Arnhold, J. and Grassberger, P. (2000) Learning driver-response relationships from synchronization patterns. Physical review. E, Statistical physics, plasmas, fluids, and related interdisciplinary topics 61, 5142–5148.

Quiroga, R. Q., Kreuz, T. and Grassberger, P. (2002) Event synchronization: a simple and fast method to measure synchronicity and time delay patterns. Physical review. E, Statistical, nonlinear, and soft matter physics 66, 041904.

Raichle, M. E. (2006) The brain’s dark energy. Science-New York Then Washington- 314, 1249.

Raichle, M. E. (2010) Two views of brain function. Trends in cognitive sciences 14, 180–190.

Raichle, M. E., MacLeod, A. M., Snyder, A. Z., Powers, W. J., Gusnard, D. A. and Shulman, G. L. (2001) A default mode of brain function. Proceedings of the National Academy of Sciences of the United States of America 98, 676–682.

Salimi-Khorshidi, G., Douaud, G., Beckmann, C. F., Glasser, M. F., Griffanti, L. and Smith, S. M. (2014) Automatic denoising of functional MRI data: combining independent component analysis and hierarchical fusion of classifiers. NeuroImage 90, 449–468.

Sanz Perl, Y., Bocaccio, H., Perez-Ipina, I., Laureys, S., Laufs, H., kringelbach, M. L., Deco, G. and Tagliazucchi, E. (2021) Non-equilibrium brain dynamics as a signature of consciousness. arXiv preprint arXiv:2012.10792.

Schaefer, A., Kong, R., Gordon, E. M., Laumann, T. O., Zuo, X. N., Holmes, A. J., Eickhoff, S. B. and Yeo, B. T. T. (2018) Local-Global Parcellation of the Human Cerebral Cortex from Intrinsic Functional Connectivity MRI. Cerebral cortex 28, 3095–3114.

Schrödinger, E. (1944) What is life? The Physical Aspect of the Living Cell. Cambridge University Press: Cambridge.

Seif, A., Hafezi, M. and Jarzynski, C. (2021) Machine learning the thermodynamic arrow of time. Nature Physics 17, 105–113.

Sejnowski, T. J. (2018) The deep learning revolution. Mit Press.

Sejnowski, T. J. (2020) The unreasonable effectiveness of deep learning in artificial intelligence. Proceedings of the National Academy of Sciences 117, 30033–30038.

Sheremet, A., Qin, Y., Kennedy, J. P., Zhou, Y. and Maurer, A. P. (2019) Wave turbulence and energy cascade in the hippocampus. Frontiers in systems neuroscience 12, 62.

Sherrington, C. (1906) The Integrative Action of the Nervous System. Yale University Press: New Haven, Connecticut.

Shirts, M. R., Bair, E., Hooker, G. and Pande, V. S. (2003) Equilibrium free energies from nonequilibrium measurements using maximum-likelihood methods. Physical review letters 91, 140601.

Silver, D., Hubert, T., Schrittwieser, J., Antonoglou, I., Lai, M., Guez, A., Lanctot, M., Sifre, L., Kumaran, D. and Graepel, T. (2018) A general reinforcement learning algorithm that masters chess, shogi, and Go through self-play. Science 362, 1140–1144.

Smith, S. M., Beckmann, C. F., Andersson, J., Auerbach, E. J., Bijsterbosch, J., Douaud, G., Duff, E., Feinberg, D. A., Griffanti, L. and Harms, M. P. (2013a) Resting-state fMRI in the human connectome project. NeuroImage 80, 144–168.

Smith, S. M., Beckmann, C. F., Andersson, J., Auerbach, E. J., Bijsterbosch, J., Douaud, G., Duff, E., Feinberg, D. A., Griffanti, L., Harms, M. P., Kelly, M., Laumann, T., Miller, K. L., Moeller, S., Petersen, S., Power, J., Salimi-Khorshidi, G., Snyder, A. Z., Vu, A. T., Woolrich, M. W., Xu, J., Yacoub, E., Ugurbil, K., Van Essen, D. C., Glasser, M. F. and Consortium, W. U.-M. H. (2013b) Resting-state fMRI in the Human Connectome Project. NeuroImage 80, 144–168.

Strogatz, S. H. (2018) Nonlinear dynamics and chaos with student solutions manual: With applications to physics, biology, chemistry, and engineering. CRC press.

van den Heuvel, M. P. and Fornito, A. (2014) Brain networks in schizophrenia. Neuropsychology review 24, 32–48.

van Vugt, B., Dagnino, B., Vartak, D., Safaai, H., Panzeri, S., Dehaene, S. and Roelfsema, P. R. (2018) The threshold for conscious report: Signal loss and response bias in visual and frontal cortex. Science 360, 537–542.

Yang, G. R. and Wang, X. J. (2020) Artificial Neural Networks for Neuroscientists: A Primer. Neuron 107, 1048–1070.

Yeo, B. T., Krienen, F. M., Sepulcre, J., Sabuncu, M. R., Lashkari, D., Hollinshead, M., Roffman, J. L., Smoller, J. W., Zollei, L., Polimeni, J. R., Fischl, B., Liu, H. and Buckner, R. L. (2011) The organization of the human cerebral cortex estimated by intrinsic functional connectivity. Journal of neurophysiology 106, 1125–1165.

Yuste, R., MacLean, J. N., Smith, J. and Lansner, A. (2005) The cortex as a central pattern generator. Nature Reviews Neuroscience 6, 477–483.

Zanin, M., Guntekin, B., Akturk, T., Hanoglu, L. and Papo, D. (2019) Time Irreversibility of Resting-State Activity in the Healthy Brain and Pathology. Frontiers in physiology 10, 1619.

Zhang, D. and Raichle, M. E. (2010) Disease and the brain’s dark energy. Nature reviews. Neurology 6, 15–28.

